# TRPM8-dependent shaking in mammals and birds

**DOI:** 10.1101/2023.12.27.573364

**Authors:** Tudor Selescu, Ramona-Andreea Bivoleanu, Mirela Iodi Carstens, Alexandra Manolache, Violeta-Maria Caragea, Debora-Elena Huțanu, Rathej Meerupally, Edward T. Wei, Earl Carstens, Katharina Zimmermann, Alexandru Babes

**Affiliations:** Department of Anatomy, Animal Physiology and Biophysics, Faculty of Biology, University of Bucharest, Bucharest 050095, Romania; Department of Neurobiology, Physiology and Behavior, College of Biological Sciences, University of California, Davis, CA 95616, USA; Department of Anesthesiology, Erlangen University Hospital, Friedrich Alexander University of Erlangen-Nuremberg (FAU), Erlangen 91054, Germany; School of Public Health, University of California, Berkeley, CA 94720, USA; Institute of Physiology and Pathophysiology, Friedrich-Alexander-University Erlangen-Nürnberg, Erlangen 91054, Germany

**Keywords:** wet dog shakes (WDS), TRPM8, cold sensing, wetness sensing, TRPM8 agonists

## Abstract

Removing water from wet fur or feathers is important for thermoregulation in warm-blooded animals. The “wet dog shake” (WDS) behavior has been largely characterized in mammals but to a much lesser extent in birds. Although it is known that TRPM8 is the main molecular transducer of low temperature in mammals, it is not clear if wetness-induced shaking in furred and feathered animals is dependent on TRPM8. Here, we show that a novel TRPM8 agonist induces WDS in rodents and, importantly, in birds, similar to the shaking behavior evoked by water-spraying. Furthermore, the WDS onset depends on TRPM8, as we show in water-sprayed mice. Overall, our results provide multiple evidence for a TRPM8 dependence of WDS behaviors in all tested species. These suggest that a convergent evolution selected similar shaking behaviors to expel water from fur and feathers, with TRPM8 being involved in wetness sensing in both mammals and birds.

## INTRODUCTION

Among the thermoregulatory behaviors necessary for limiting heat loss and surviving in wet-cold conditions, one of the most spectacular is the shaking behavior mammals and birds use to efficiently expel water from their coat. This behavior, called in mammals “wet-dog shake” (WDS), is a shudder motion consisting in vigorous and rapid rotations of the head and trunk around the spinal axis, with frequencies depending on body mass (1). While the mammalian WDS behavior has been the subject of biomechanics and pharmacological research (1–6), the analogous behavior in birds received little attention (7) and it was not investigated so far to what extent it can be triggered pharmacologically. By bringing together results from pharmacology, genetics and naturalistic stimulation, our study aimed to obtain a deeper understanding of the factors triggering WDS-like behaviors in mammals and birds.

Using WDS as a proxy can also be a useful approach to study wetness sensing. By modulating the peripheral nervous system-dependent component of WDS, wetness sensing can be studied objectively and quantitatively in animal models. This allows genetic and pharmacological manipulations, which would constitute an advantage compared to human psychophysics (8).

Icilin-induced WDS and jumping were previously shown in mice to be dependent on the cold and menthol receptor – the Transient Receptor Potential subfamily M (melastatin) member 8 (TRPM8) cation channel, as TRPM8 knock-out (*Trpm8-/-*) mice do not display icilin-induced WDS and jumping (9, 10). In rats, the dependency on TRPM8 for icilin-induced WDS was demonstrated using TRPM8 antagonists (11–13). Nevertheless, the relationship between the pharmacologically induced behavior and natural shaking of wet animals was not investigated. The question whether the shaking of rodents, when sprayed with cold water, is dependent on TRPM8, is therefore still unanswered.

Thus far, the study of TRPM8 dependent shaking in rodents relied exclusively on the use of icilin. Because avian TRPM8 is insensitive to icilin (14), the investigation of TRPM8 involvement in a putative bird shaking behavior has been hampered. A WDS-like shaking behavior in birds has been experimentally characterized only in one particular case (Anna’s hummingbirds, *Calypte anna*, perched and during flight (7). Available anecdotal evidence shows that chickens and other birds do shake to expel water from their plumage. We proceeded to confirm this with controlled investigations under laboratory conditions.

Thus, given the incomplete understanding of the sensory mechanisms that trigger shaking in wet mammals and birds, we embarked on a study to explore in detail the role played by the cold sensor TRPM8 in these behaviors. For this purpose, we used a novel TRPM8 agonist (15, 16) to overcome the limitations of icilin in activating avian TRPM8. We then set out to find what role TRPM8 plays in wetness-dependent WDS of mice under naturalistic stimulation. Our study will allow a better understanding of how shaking to remove water is triggered and how it evolved.

## RESULTS

### C-1 is a potent agonist of avian TRPM8

To find a TRPM8 agonist able to elicit WDS-like behaviors in both birds and mammals, we started with pharmacological assays in heterologously expressed TRPM8 orthologs from chicken (cTRPM8), rat (rTRPM8) and human (hTRPM8). We considered a novel TRPM8 agonist, 1-diisopropylphosphorylheptane (DIPA-1-7), also known as Cryosim-1 (C-1), due to its good water solubility, which is advantageous for *in vivo* administration. C-1 belongs to a new class of “cooling compounds’’, called diisopropylphosphinoylalkanes (DIPA), structurally distinct from menthol and its derivatives (e.g., WS-12), and from icilin (Fig. 1A, upper part). Three TRPM8 agonists (C-1, WS-12 and icilin) were tested at equal concentrations (10 µM) under temperature control of the perfusate, using calcium imaging of HEK293T cells transfected with c/r/hTRPM8. As expected, icilin activated only the mammalian orthologs, while C-1 and WS-12 activated all orthologs (Fig. 1B).

**Fig. 1.**
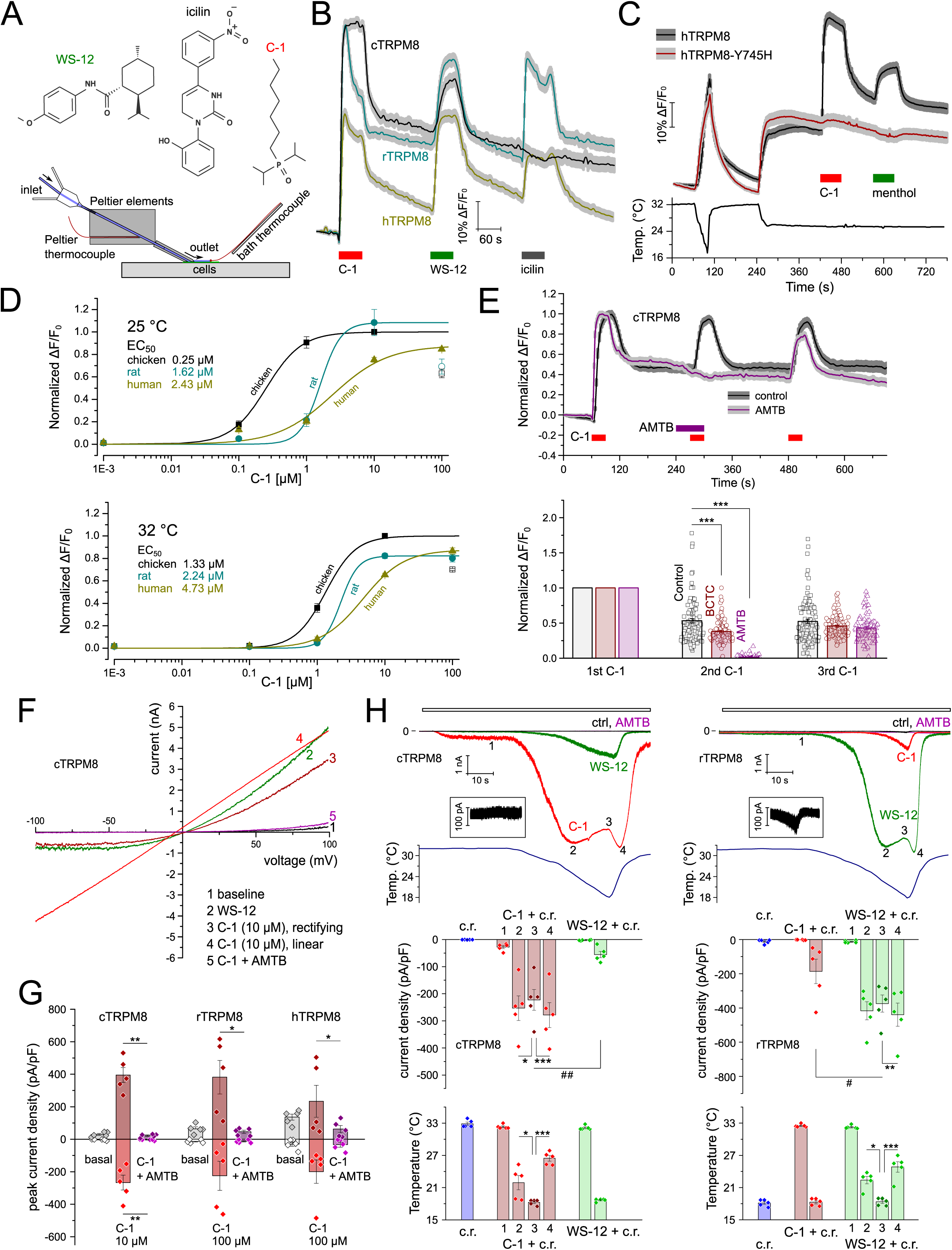
The main properties of the novel agonist C-1 on recombinant TRPM8 from mammals and birds. (A) Top: The structures of synthetic TRPM8 agonists WS-12, icilin, and C-1 showing they belong to distinct classes of cooling-mimetic compounds. Bottom: the schematic diagram of the rapid temperature control system using Peltier elements and feedback from two thermocouples. (B) Calcium imaging traces (ΔF/F_0_) of HEK293T cells expressing one of c/r/hTRPM8, stimulated with the three agonists: C-1, WS-12, icilin, all at 10 µM (129/110/93 cells for c/r/hTRPM8, respectively). Only cTRPM8 showed a sustained response to C-1, reflected in the larger area under the ΔF/F_0_ curve (AUC) (Fig. S1A). (C) The menthol-insensitive Y745H mutant of hTRPM8 is also insensitive to C-1 (100 µM). Cooling from 32 to 17 °C, and then from 32 to 25 °C, functionally confirmed the expression of hTRPM8-Y745H. Menthol (100 µM) was applied at the end of the experiments to confirm the differences in menthol sensitivity (averaged traces from 114 cells expressing hTRPM8 and 126 cells expressing hTRPM8-Y745H). (D) Concentration-response curves for c/r/hTRPM8 at 25 and 32 °C. Each ΔF/F_0_ data point for C-1 (0.1-100 µM) is averaged from 3-4 experiments, with 17-74 cells per experiment; the average ΔF/F_0_ for each ortholog was normalized to the response of cTRPM8 to 10 µM C-1. Data represented by filled symbols were fitted with Hill equations; empty symbols show desensitized responses at 100 µM C-1. The values at the left side of each graph represent the EC_50_ values, showing that cTRPM8 is the most sensitive ortholog to C-1, while hTRPM8 the least sensitive. (E) Top: Mean ± SEM traces (ΔF/F_0_) for control and antagonist experiments with cTRPM8 stimulated three times by C-1 (10 µM, 30 s each). The TRPM8 antagonist AMTB (1 µM) was applied before and during the second C-1 challenge. The traces were normalized to the response to the first C-1 challenge (45/85 cells from control/AMTB experiments). Bottom: the normalized means of maximal ΔF/F_0_ from experiments with AMTB and another TRPM8 antagonist, BCTC (10 µM). AMTB completely and reversibly inhibited the calcium transient elicited by C-1 (***, *p*<0.001, *t*-test, paired; 98/85/98 cells from control/AMTB/BCTC experiments). AMTB and BCTC also inhibited r/hTRPM8 (Fig. S1B). (F) Representative examples of I-V relationships for cTRPM8 expressed in HEK293T cells when superfused with WS-12 (10 µM), C-1 (10 µM) and C-1 plus AMTB (1 µM). The temperature was kept at 25 °C. Trace ‘3’ represents an outwardly rectifying I-V curve (before the maximum current was reached) and ‘4’ a linear relationship, recorded during peak current. Similar I-V relationships were found for rTRPM8 and hTRPM8 (Fig. S1C). (G) Data pooled from experiments as in F. Inward and outward currents elicited by C-1 (10 µM in cTRPM8 and 100 µM in rTRPM8 and hTRPM8) were fully inhibited by AMTB (**, *p*<0.01; *, *p*<0.05, *t*-test, paired, n=5 for each ortholog). C-1 (100 µM) elicited significantly larger currents than WS-12 (10 µM) in cTRPM8 (Fig. S1D). (H) (**Top**): Representative inward currents recorded at –80 mV in the HEK293T cells expressing cTRPM8 or rTRPM8 during cooling ramps (c.r.) alone (from 32-33 to 18 °C) and then together with C-1 (10 µM), WS-12 (10 µM), and AMTB (1 µM). The maximum current recorded before the beginning of the cooling ramp is marked with ‘1’. Currents recorded while the cooling ramp and a potent agonist were concomitantly applied show a local minimum (marked with ‘3’, between the two maxima at ‘2’ and ‘4’). cTRPM8 displayed a current dip to C-1, while rTRPM8 to WS-12. The same was confirmed in hTRPM8 (Fig. S1E). Insets show at higher resolution the peak currents elicited by cooling alone. (**Bottom**) Data pooled from experiments as above, showing the average current densities and corresponding temperatures at which the peak currents were recorded. In cTRPM8, C-1 during cooling elicited larger currents than WS-12 during cooling, while the reverse was true for rTRPM8 (for hTRPM8, see Fig. S1E; ***, *p*<0.001; **, *p*<0.01; *, *p*<0.05; one-way repeated measures ANOVA, n=5 for each ortholog; ##, *p*<0.01; #, *p*<0.05, repeated measures two-way ANOVA followed by Tukey’s post-hoc test, n=5 for each ortholog). All traces and columns represent means ± SEM. **Figure 1 – Source data 1:** current densities and bath temperature values used to generate the bar plots in Figure 1G, H.

Tyrosine 745 is known to play an important role in the interaction of menthol with its binding pocket: the Y745H mutant of hTRPM8 is menthol-insensitive but nonetheless cold-sensitive (17, 18).

A cooling ramp and a cooling step elicited similar responses in both WT and Y745H hTRPM8, while C-1 and menthol induced large calcium transients only in WT hTRPM8. This shows that C-1 binds to the channel in the same manner as most other TRPM8 agonists (17) (Fig. 1C).

To compare the C-1 sensitivity of the three orthologs, the ΔF/F_0_ responses to a range of concentrations (0.1-100 µM C-1) were normalized to the ΔF/F_0_ response of cTRPM8 to 10 µM C-1 and fitted with Hill equations. As expected for TRPM8, the EC_50_ values for each species were lower at 25 °C than at 32 °C. At both temperatures, cTRPM8 had the lowest EC_50_, while hTRPM8 the highest (Fig. 1D, Table S1).

The selective TRPM8 antagonist AMTB abolished the calcium transients elicited by C-1 in all three orthologs (Fig. 1E, Fig. S1B). The inhibition was rapidly reversible only for cTRPM8. BCTC, a well-characterized TRPV1 and TRPM8 antagonist (18), had a lower inhibitory effect compared to AMTB, particularly on cTRPM8 (Fig. 1E).

The properties of heterologously expressed c/r/hTRPM8 were investigated using whole-cell patch clamp current recordings under calcium-free conditions. All three orthologs showed current-voltage relationships with TRP-characteristic outward rectification, which often became linear when the channels were maximally activated (Fig. 1F, Fig. S1C). AMTB inhibited the outward and inward whole-cell currents activated by C-1 (Fig. 1G). When comparing the currents elicited by C-1 (100 µM) and WS-12 (10 µM) in the same cells, cTRPM8 exhibited larger currents to C-1 than to WS-12 (Fig. S1D).

In further experiments, currents were recorded at –80 mV, while the temperature was decreased from ∼32 to ∼18 °C (cooling ramp of ∼37 s) during stimulation by agonists and antagonists. Cooling alone evoked small but measurable inward currents in r/hTRPM8, but not in cTRPM8 (Fig. 1H, Fig. S1E), in line with results of previous investigations (19). As expected, in all orthologs, C-1 and WS-12 enhanced the currents elicited by the cooling ramp, while AMTB abolished them (Fig. 1H, Fig. S1E). The maximum 1/Q_10_ coefficients for the temperature-dependent inward current increase in the presence of agonists were between 139 and 820, depending on the ortholog – agonist pairing (Table S1). Interestingly, the current decreased when approaching the minimum temperature, during exposure to C-1 for cTRPM8, and to WS-12 for r/hTRPM8, producing a “W”-shaped current. The local current minimum coincided with the temperature minimum during the cooling ramp. The Q_10_ coefficients for the temperature-dependent current decrease ranged from 1.2 to 1.3 for all TRPM8 orthologs (Table S1). This shows that TRPM8 exhibits diffusion-limited conductivity at its maximal opening probability (Fig. 1H, Fig. S1E). It also shows that C-1 is an efficacious agonist of cTRPM8 at temperatures below ∼22.5 °C.

### Native TRPM8 is selectively activated by C-1 in cultured DRG neurons from rodents and chickens

The sensitivity of native TRPM8 to C-1 was investigated in cultured sensory DRG neurons from chicken, rat and mouse (WT and *Trpm8-/-* mice). In rat DRG cultures, microscopic fields containing neurons likely to express TRPM8 were selected by a slight cooling step (from 32 to 25 °C). Their sensitivity to the well-established TRPM8 agonist, WS-12 (5 µM), and to C-1 (10 or 100 µM) was then tested at a constant temperature of 25 °C (Fig. 2A, top). The large overlap in sensitivity to C-1 (at both concentrations used) and to WS-12, strongly suggests that TRPM8 was selectively activated by C-1 (Fig. 2A, bottom). The mild temperature step revealed the most cold-sensitive neurons, which were also, to a large extent, C-1 sensitive (Fig. S2).

**Fig. 2.**
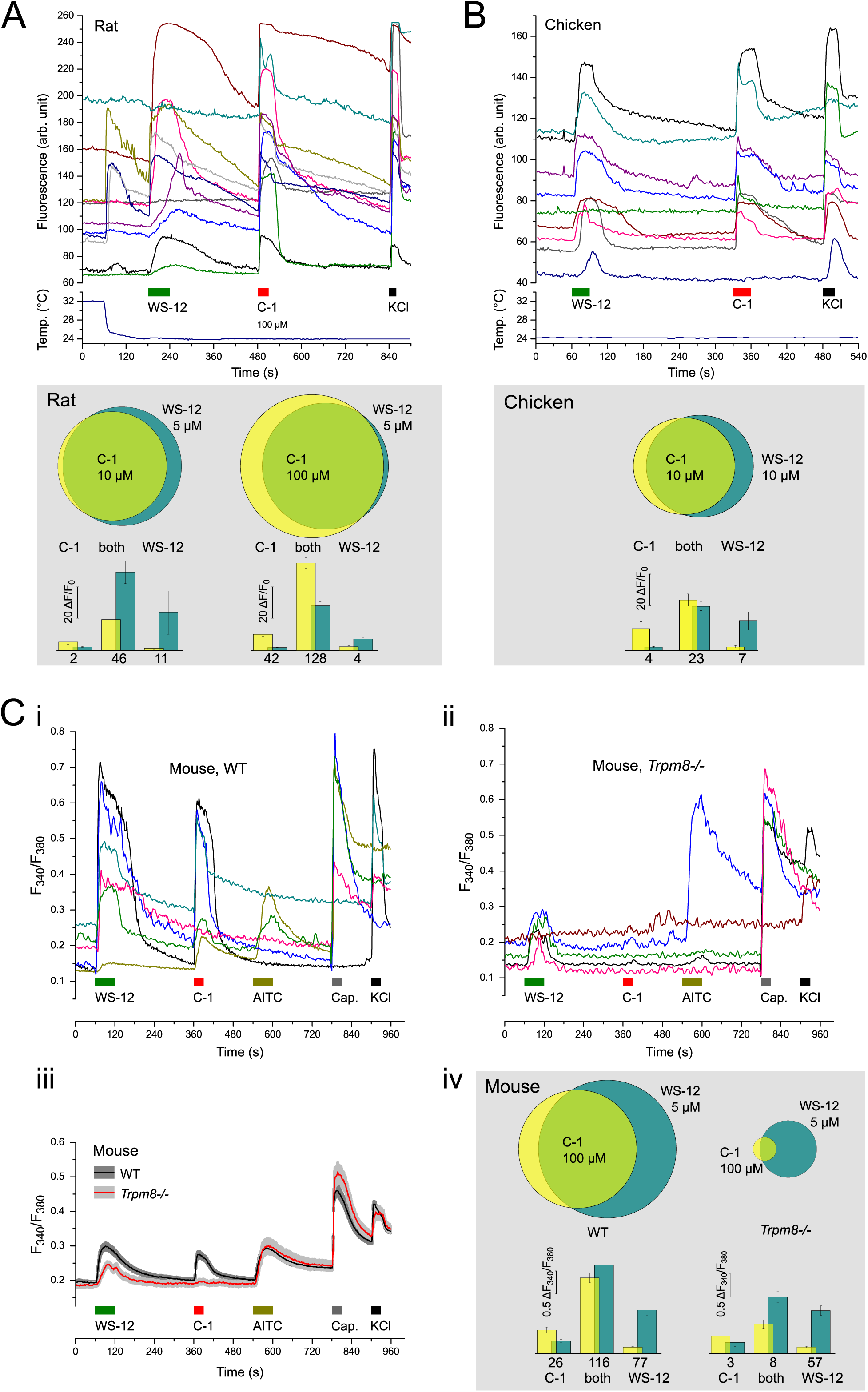
Native TRPM8 from rat, mouse, and chicken DRG neurons is selectively activated by C-1. (A) (**Top**) representative examples of fluorescence traces in cultured DRG neurons from rat. The mild cooling step (from 32 to 25 °C) was used to preselect neurons that would likely express TRPM8 and was followed by the established TRPM8 agonist, WS-12, and the novel agonist, C-1. A high potassium concentration (50 mM KCl) was used at the end of the experiments to identify viable neurons. (**Bottom**) Venn diagrams illustrating the overlap between the neurons responding to WS-12 (5 µM) and C-1 (10 or 100 µM). Experiments were performed on 456 neurons (10 µM C-1) and on 970 neurons (100 µM C-1). The statistics at the bottom show the average peak amplitudes (ΔF/F_0_) to C-1 and WS-12, as well as the neuron numbers for each of the three populations: neurons passing the threshold for C-1 only, for both agonists, and for WS-12 only. (B) (**Top**) similar to A, in chicken DRG neurons, exposed to WS-12 (10 µM), C-1 (10 µM) and high potassium (50 mM KCl) at constant temperature (24-25 °C). (**Bottom**) Venn diagram illustrating the overlap in the responses to WS-12 and C-1. Experiments were performed on 369 neurons. The statistics at the bottom show the average peak amplitudes (ΔF/F_0_) and neuron numbers for each of the three populations. (C) Representative ratiometric imaging traces from wild-type C57BL/6 WT (**i**) and *Trpm8-/-* mice (**ii**). To characterize the neurons, WS-12 (5 µM), C-1 (100 µM), AITC (100 µM), capsaicin (1 µM) and KCl (50 mM) were superfused at constant temperature (∼25 °C). (**iii**) The average F_340_/F_380_ traces (± SEM) of all neurons responding to either WS-12 or C-1, from WT (n=219 neurons) and *Trpm8-/-* (n=68 neurons) mice. (**iv**) Venn diagrams displaying the overlap of responses to WS-12 and C-1 in neurons from both WT and *Trpm8-/-*. Experiments were performed on 1167 neurons from WT mice and 1245 neurons from *Trpm8-/-* mice. The statistics at the bottom illustrate the average peak amplitudes (ΔF_340_/F_380_) and neuron numbers for each of the three populations. The scaling of the Venn diagrams reflects the ratios of the percentages of responding cells from the total neuronal populations investigated. **Figure 2 – Source data 1:** individual neuronal amplitudes (ΔF_340_/F_380_) of the responses to WS-12 and C-1 in DRG neurons from WT and *Trpm8-/*-mice, used to generate the Venn diagrams and average amplitudes in Figure 2C iv.

In cultured chicken DRG neurons, WS-12 (10 µM) was used to select microscopic fields rich in TRPM8-expressing neurons before stimulation with C-1 (10 µM). The temperature stimulus was not reliable for selecting TRPM8-expressing neurons, because cooling also elicited numerous TRPM8-independent calcium transients in chicken sensory neurons, as shown previously (20).

Therefore, the recordings in chicken DRG neurons were performed at a constant temperature of 25 °C (Fig. 2B, top). Considering the sensitivity of recombinant cTRPM8, we chose a concentration of 10 µM, for both C-1 and WS-12, in chicken DRG neuron assays. Again, a substantial overlap was observed between C-1 and WS-12 sensitivities (Fig. 2B, bottom).

To unambiguously determine the role of TRPM8 in eliciting calcium transients in response to C-1 and WS-12, we used DRG neurons from WT and *Trpm8-/-* mice and ratiometric calcium imaging with Fura-2 at ∼25 °C. Cells were sequentially stimulated with TRPM8, TRPA1, and TRPV1 agonists, as shown in Fig. 2C.

The averaged ratiometric signal of all neurons responding to either WS-12 or C-1 in WT mice (219 neurons sensitive to WS-12 and/or C-1 from a total of 1167) and *Trpm8-/-* mice (68 neurons sensitive to WS-12 and/or C-1 from a total of 1245), allowed us to visually discriminate the effect of the absence of *Trpm8-/-* on responses to the two agonists (Fig. 2C, iii). We noticed that the WS-12 responses of *Trpm8-/-* mice were still preserved (albeit fewer and with lower amplitude compared to WT). These responses to WS-12 in *Trpm8-/-* neurons were well synchronized as revealed by the shape of the average response (Fig. 2C, iii), suggesting a single TRPM8-independent responsible mechanism. In *Trpm8-/-* mice, responses to WS-12 alone were represented by a substantial population of neurons, 57 from 1245 neurons (4.58%).

Moreover, the average ΔF_340_/F_380_ amplitudes of neurons responding only to WS-12 were the same in neurons from WT and *Trpm8-/-* mice (Fig. 2C, iv). All the above suggest that, in mouse DRG neurons, C-1 has a higher selectivity towards TRPM8 compared to WS-12. Nevertheless, the total number of neurons responding to either C-1 or WS-12 was significantly lower in *Trpm8-/-* than in WT (chi-square goodness-of-fit test, *p*<0.0001): 11 from 1245 versus 142 from 1167 (for C-1), and 65 from 1245 versus 193 from 1167 (for WS-12, Fig. 2C, iv).

### C-1, icilin, and water-spraying evoke similar WDS in rats

The property of icilin to induce WDS in a variety of mammals was discovered long before the cloning and characterization of TRPM8. Most of the early WDS experiments with icilin were carried out in rats (21–23).

The generation of *Trpm8-/-* mice helped showing that WDS and jumping behaviors triggered by icilin are completely dependent on TRPM8 (9, 10). The icilin WDS assay in rats became a standard method that allowed *in vivo* validation of TRPM8 antagonists (11–13).

In rats, a complete WDS event begins with a rotation of the head and continues towards the tail, often with the fur covering distant body segments rotating in opposite directions (Fig. 3A, Movie S1). To compare the effect of C-1 with the known WDS response to icilin, Wistar rats were injected intraperitoneally (i.p.) with either 33 mg/kg C-1 or 1 mg/kg icilin. Both treatments elicited robust WDS, with a similar number of bouts, lasting up to 1 h. In all rats injected with either drug, WDS bouts were frequently followed by grooming (Fig. 3B). WDS bouts were counted both over 5-min intervals (Fig. 3C) and for the entire session (30 min, Fig. 3D). Neither substance elicited writhing behavior. Injections with vehicles had no effect on the measured behavioral parameters (Fig. 3C). WS-12, at a dose of up to 33 mg/kg, was also injected i.p. into 4 rats without eliciting any WDS (data not shown).

**Fig. 3.**
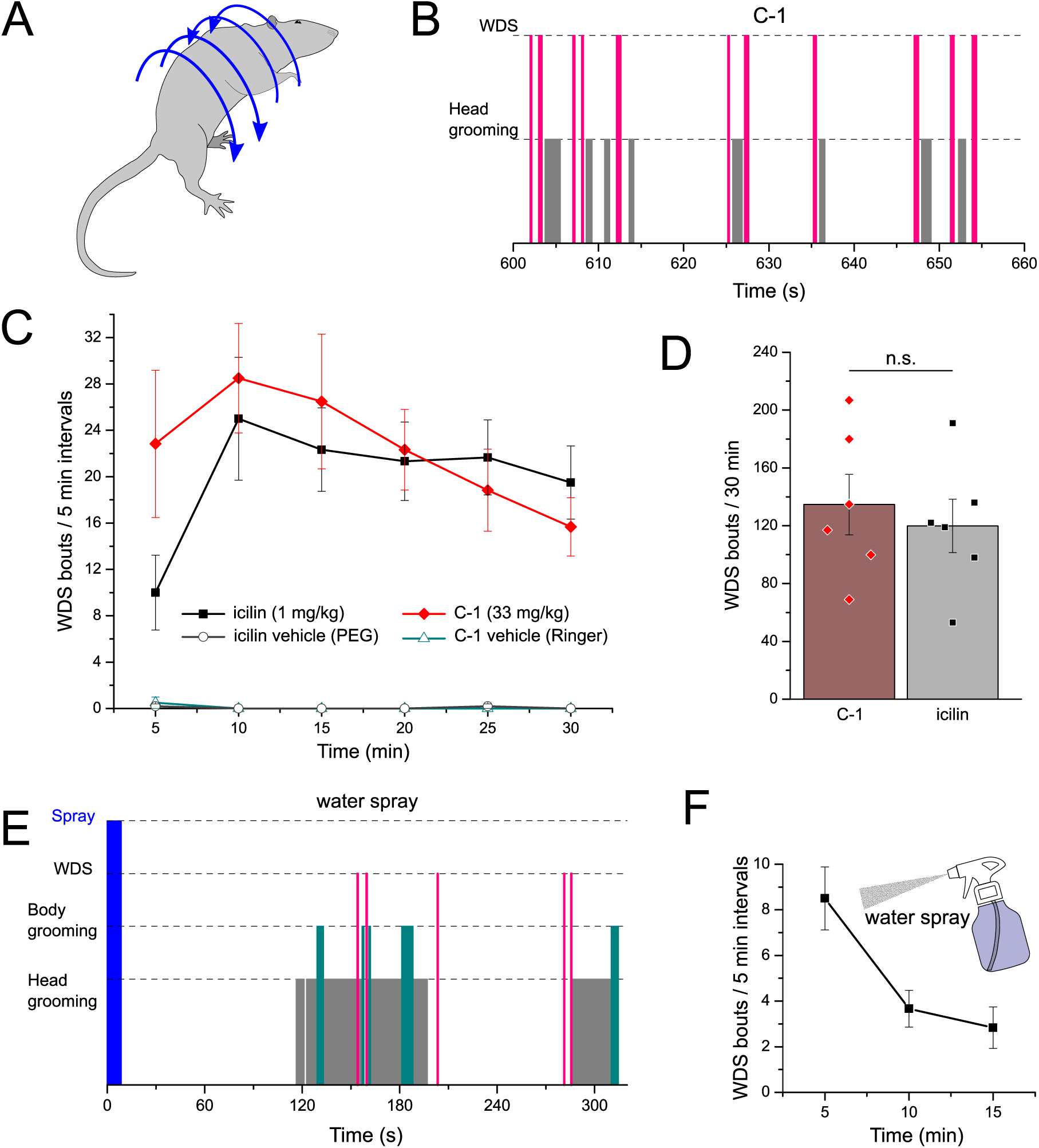
In rats, C-1 and icilin elicit comparable WDS behaviors, stronger than water spraying. (A) The typical rotational movements during WDS in a rat. During a vigorous bout, the head rotates in the opposite direction relative to the fur on the rear of the body. (B) A representative ethogram (1 min), starting at 10 min after C-1 was administered. Grooming often followed the WDS bouts (examples shown in Movie S1). This was typical for all recorded animals. (C) The time-course of WDS bouts in rats that received i.p. C-1 (33 mg/kg) or icilin (1 mg/kg), and for each, their respective vehicles (Ringer solution and PEG). The data points represent the number of WDS bouts counted over 5 min intervals (n=6 for each experiment). (D) The total number of WDS bouts from experiments in C. C-1 and icilin elicited similar numbers of WDS bouts over 30 min (n.s., *p*>0.05, *t*-test, unpaired, n=6). (E) A representative ethogram of a rat sprayed with water at room temperature (∼23 °C, 20 sprays with 0.9 ml/spray, in quick succession), showing the behaviors recorded during 5 min from spraying. (F) The time-course of WDS bouts in sprayed rats counted over 5 min intervals (n=6). All points and columns represent means ± SEM.

To investigate the rats’ natural response to water droplets, similar to rain, the animals were sprayed 20 times in quick succession with a pump bottle delivering 0.9 ml of water per spray at room temperature (∼23 °C). Rats often approached the spray source, indicating that the spraying of water was not perceived as an aversive stimulus. The spray-evoked WDS movement was nearly identical to the chemically-triggered one. The number of WDS bouts, counted up to 15 min after spraying, was lower than in the rats receiving i.p. C-1 or icilin (Fig. 3E, F).

### TRPM8 is required for C-1-triggered WDS in mice

To test whether C-1-triggered WDS are dependent on TRPM8, the behavior of WT and *Trpm8-/-* mice was compared. Wild-type and *Trpm8-/-* mice were injected i.p. with 33 mg/kg C-1 and their behavior was recorded for 30 min. As we found no statistically significant sex differences, mice of both sexes were pooled. Following the injection, a crouched posture and an early-onset (<1 min) high-frequency grooming interspersed with WDS bouts were detected in WT mice (Fig. 4A-C, Movie S2). In contrast, WDS bouts were almost absent in *Trpm8-/-* mice during the entire 30 min of the experiment while the grooming duration was significantly shorter (Fig. 4C, D).

**Fig. 4.**
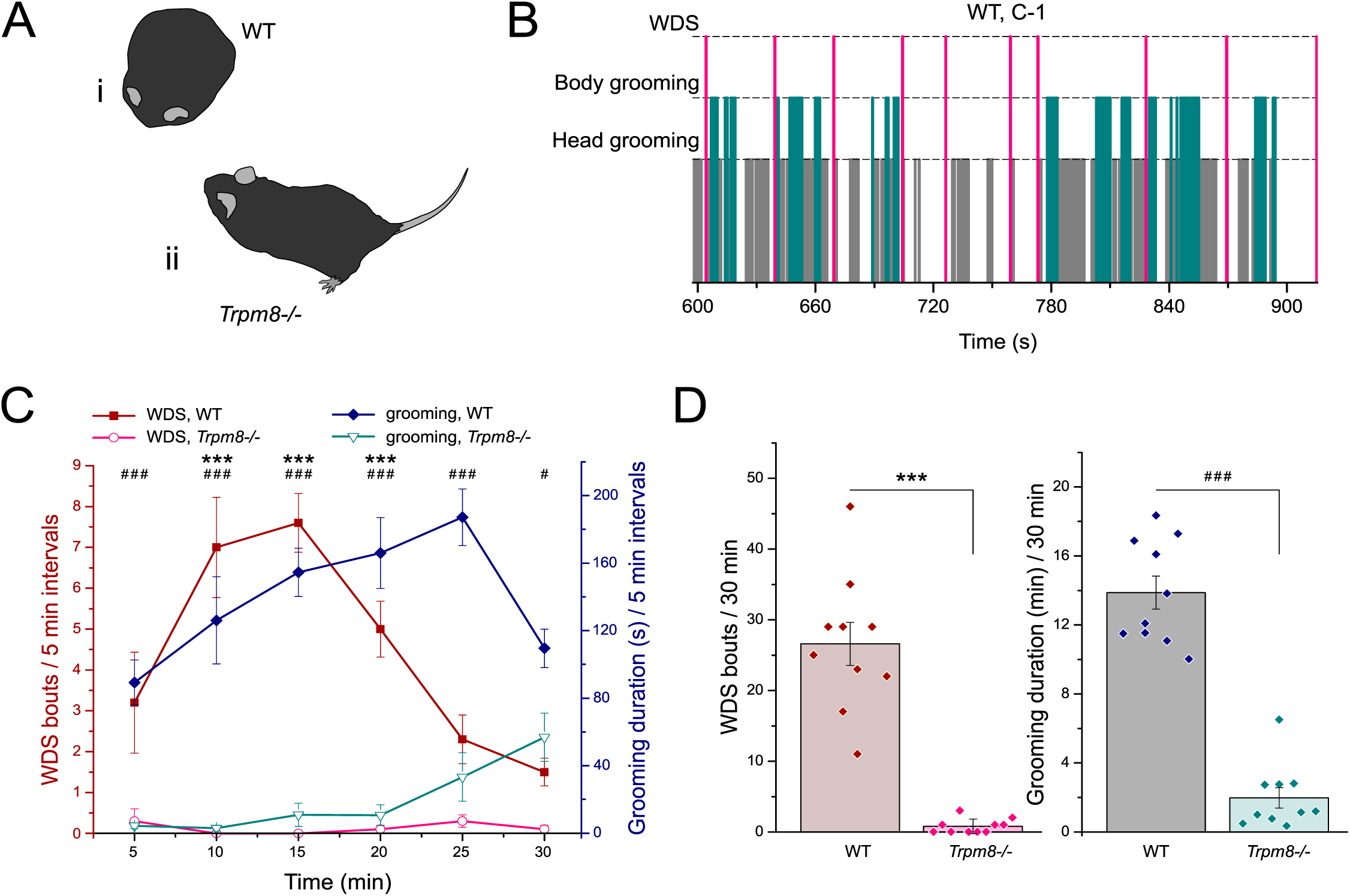
TRPM8 is required for C-1-elicited WDS and grooming in mice. (A) Typical postures of WT (**i**) and *Trpm8-/-* (**ii**) mice after receiving i.p. C-1 (33 mg/kg): WT mice adopted a crouched posture most of the time, while *Trpm8-/-* mice displayed a normal posture (examples shown in Movie S2). (B) A representative ethogram (5 min) of a WT mouse behavior, starting at min 10 after C-1 injection. (C) The time-course of WDS bouts (left Y-axis) and grooming duration (right Y-axis) in WT and *Trpm8-/-* mice after receiving i.p. C-1. Data points represent the average number of WDS bouts and grooming duration counted over 5 min intervals (for WDS bouts: ***, *p*<0.001; for grooming: ###, *p*<0.001; #, *p*<0.05, repeated measures two-way ANOVA followed by Tukey’s post-hoc test, n=10 for each genotype). (D) Total WDS bouts (left) and grooming duration (right) from experiments in C, counted over 30 min. In WT mice, i.p. C-1 elicited significantly more WDS bouts and grooming than in *Trpm8-/-* mice (for WDS bouts: ***, *p*<0.001; for grooming: ###, *p*<0.001, *t*-test, unpaired, n=10 for each genotype). All points and columns represent means ± SEM.

### TRPM8 is required for temperature-dependent WDS latency in water-sprayed mice

The response of mice to water spray stimuli was investigated in both WT and *Trpm8-/-* mice, using water at two temperatures. The aim of these experiments was to separate the cooling-induced effects from mechanically-induced ones, since, during rain, an animal is exposed to both a decrease of fur temperature and the impact of droplets. Preliminary observations showed that C57BL/6 mice shake in response to mechanical stimuli alone (dust sprinkling on their coat, Movie S3), thus we did not expect a complete absence of shaking behaviors in *Trpm8-/-* mice, or in WT mice sprayed with warm-neutral water, but a temperature modulation of WDS bouts number, and especially of the WDS latency. Therefore, one target temperature was chosen to be close to the initial temperature measured on the animals’ dorsal fur (∼32 °C, hereafter referred to as “warm” spraying), so that we would only capture the effects of the mechanical components immediately after spraying. The other target temperature was chosen to reflect significant cooling, without entering the painfully cold range (below 12 °C, (24), hereafter referred to as “cold” spraying). Spraying consisted of 4 rapid successive actuations of a pump bottle, each delivering ∼0.85 ml of distilled water (Movie S4). The measured dorsal fur temperatures were 32.0 ± 2.4 °C (warm, n=26) and 14.5 ± 1.6 °C (cold, n=26), while the average temperatures inside the spray bottle were ∼58 and ∼4 °C, respectively (Fig. 5A, B, D).

**Fig. 5.**
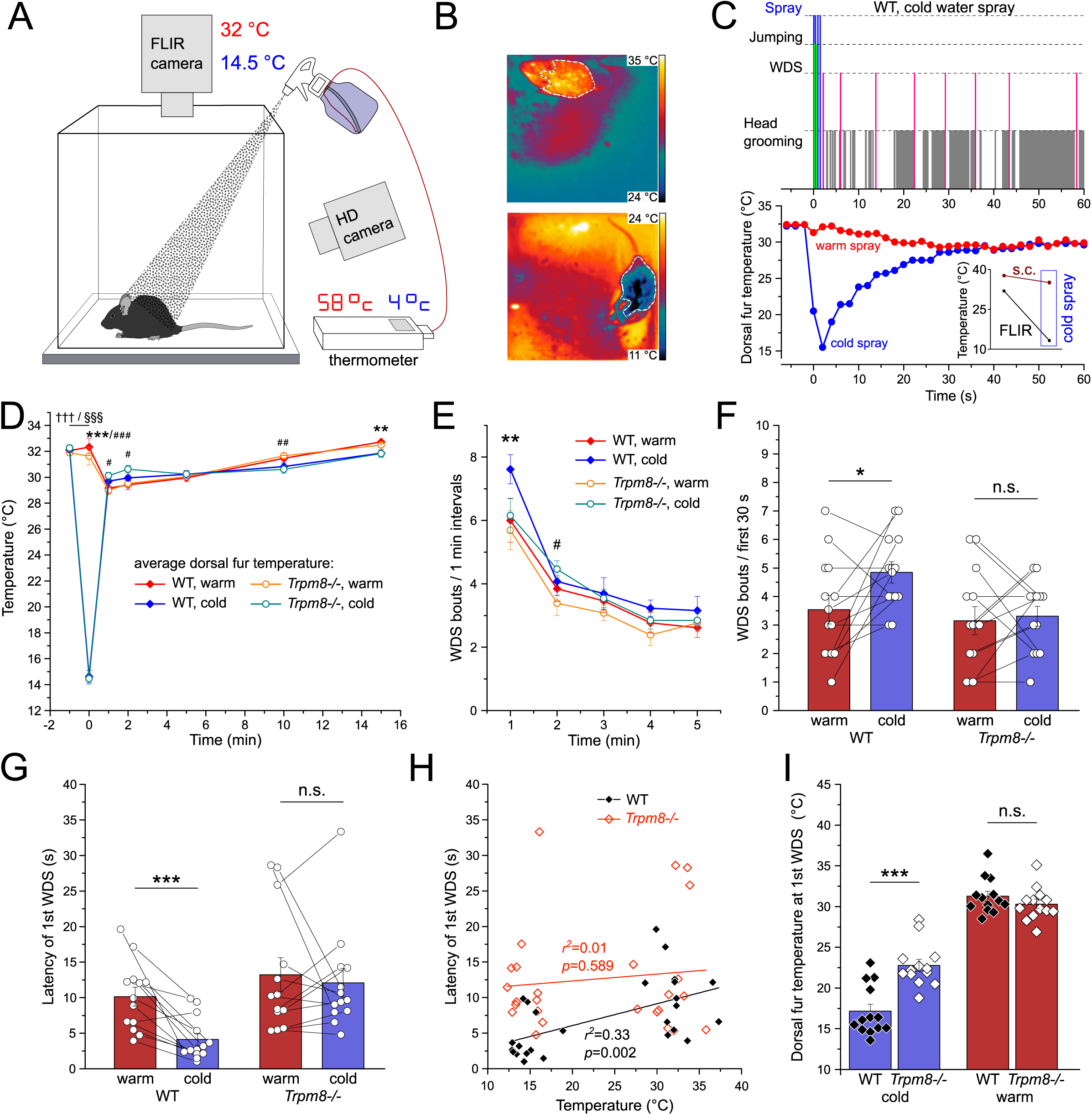
TRPM8 is required for short latency shaking in cold-water-sprayed mice. (A) The system used for recording the behavior and dorsal fur temperature of water-sprayed mice. Each WT and *Trpm8-/-* mouse was sprayed (4 sprays with ∼0.85 ml/spray, in quick succession) in separate experiments with warm and cold water (Movie S4). The measured dorsal fur temperatures were 32.0 ± 2.4 °C (warm, n=26) and 14.5 ± 1.6 °C (cold, n=26). The corresponding water temperatures inside the spraying bottles necessary to reach the above temperatures on the mouse dorsal fur, were ∼58 and ∼4 °C, respectively. (B) Color-coded radiometric images recorded with the FLIR camera when a mouse was sprayed with warm (top) and cold water (bottom). The white polygonal contours demarcate the areas used for measuring the average temperature of the mice’s dorsal fur after spraying. (C) A composite figure showing a representative ethogram of a WT mouse sprayed with cold water (first minute after spraying, upper part), synchronized with the FLIR measured temperature of the dorsal fur (“cold spray”, lower part). The “warm spray” trace was added for comparison and illustrates the temperature variation of the same mouse when sprayed with warm water. Inset: in a separate group of mice, s.c. implanted with temperature transponders, cold spraying decreased the dorsal fur temperature (FLIR measurements) from a 31.9 ± 0.2 °C baseline to 13.2 ± 0.5 °C (n=5), while the back s.c. temperature decreased from a 37.7 ± 0.2 °C baseline to 35.1 ± 0.7 °C (n=5). (D) The time-course of mice dorsal fur temperature in WT and *Trpm8-/-* mice before and after warm and cold spraying. The largest temperature differences were measured immediately after spraying (dorsal fur difference 17.4 ± 0.5 °C). The differences between the dorsal fur temperatures resulting from warm and cold spraying were analyzed for each genotype (comparisons between warm and cold spraying at the same time points: *** and ###, *p*<0.001; ** and ##, *p*<0.01; #, *p*<0.05, repeated measures two-way ANOVA followed by Tukey’s post-hoc test, n=12 for WT (*) and n=13 for *Trpm8-/-* (#); comparison between basal temperature and immediately after cold spraying: ††† (WT) and §§§ (*Trpm8-/-*), *p*<0.001, *t*-test, paired, n=13 for each genotype). (E) WDS bouts counted in 1 min intervals during the first 5 min after spraying. In WT mice, cold spraying evoked significantly more WDS bouts than warm spraying in the same mice, during the first minute (**, *p*<0.01; #, p<0.05; repeated measures two-way ANOVA followed by Tukey’s post-hoc test, n=13 for both WT (*) and *Trpm8-/-* (#)). (F) The number of WDS bouts immediately after the spray. *Trpm8-/-*, but not WT mice, performed a lower number of cold-induced WDS bouts in the first 30 s of the test (*, *p*<0.05; n.s., *p*>0.05, *t*-test, paired, n=13 from each genotype). (G) The latency of the first WDS in WT and *Trpm8-/-* mice in spraying experiments. WT mice had significantly shorter latencies of WDS in response to cold than warm spraying, while the difference was not statistically significant in *Trpm8-/-* mice (***, *p*<0.001; n.s., *p*>0.05, *t*-test, paired, n=13 from each genotype). (H) First WDS latency displayed against individual dorsal fur temperature after spraying in WT and *Trpm8-/-* mice. The dorsal fur temperature of each mouse after cold and warm water-spraying is shown on the X-axis. Simple linear regressions show that the WT mice have a temperature-dependent shaking latency. The coefficients of determination (*r^2^*) and *p-*values are displayed in the graph. (I) The dorsal fur temperature measured immediately before the first WDS was significantly different between WT and *Trpm8-/-* mice for cold spraying, but not for warm spraying (***, *p*<0.001; n.s., *p*>0.05, *t*-test, unpaired, n=13 for each condition). **Figure 5 – Source data 1:** dorsal fur temperature values, number of WDS bouts and WDS latencies for Figure 5D-I.

Post-spray behavior consisted of WDS and grooming, while some animals also showed a startle-like response, by jumping off the ground with all four paws in response to single sprays. In a typical response, the frequency of WDS bouts decreased, while the grooming duration increased, as the fur temperature recovered after spraying (Fig. 5C). The dorsal fur temperature, measured immediately after cold spraying, was significantly lower than the baseline temperature (*p*<0.001, n=13 for each genotype, Fig. 5D) and it quickly recovered to baseline (in less than 1 min, Fig. 5C, *t*-testD). There were no significant differences between the dorsal fur basal temperature and the fur temperature measured immediately after warm spraying, as we intended (*p*>0.05, paired-sample, n=13 for each genotype). The dorsal fur temperature difference between warm and cold spraying was analyzed immediately and at 1, 2, 5, 10, and 15 min after spraying. As expected, the largest significant difference was found immediately after spraying (Fig. 5D).

Because the fur temperature differences between warm and cold spraying decreased greatly over time, we counted WDS bouts in 1-min intervals during the first 5 min from spraying. The number of WDS bouts decreased over time in all conditions (Fig. 5E). During the first minute, WT mice sprayed with cold water showed a significantly greater number of WDS bouts than after warm water-spraying (Fig. 5E). In contrast, the WDS bout difference in the first minute was not significant in *Trpm8-/-* mice. In these mice there was a significant difference only for the interval between the first and second minute from spraying (Fig. 5E). When the interval for counting WDS was reduced to the first 30 s after spraying, a similar difference as at the first minute was found between genotypes (Fig. 5F).

Considering the short-lasting effect of cooling the fur after spraying, we focused our analysis on the minimum duration necessary to record the WDS behavior, which is the latency of the first WDS. This was measured from the onset of spraying. WT mice had an average latency of the first WDS to cold spraying of 4.13 ± 0.84 s, less than half of the latency to warm spraying, 10.14 ± 1.32 s (*p*<0.001, n=13). *Trpm8-/-* mice showed no significant differences in WDS latency between warm and cold spraying, showing their WDS onset was not sensitive to fur temperature (Fig. 5G).

To analyze the relationship between individual spray temperatures and WDS latencies in WT and *Trpm8-/-* mice, simple linear regressions were fitted to the data (Fig. 5H). A significant relationship was found for WT mice (*p*<0.01), with a positive correlation between latency and fur temperature (*r^2^*=0.33).

Likely as a consequence of the different WDS latencies, combined with the rapid recovery of the fur temperature towards baseline, the fur temperature measured immediately before the first WDS evoked by cold spraying was significantly lower in WT mice than in *Trpm8-/-* mice (17.2 ± 0.9 versus 22.8 ± 0.7 °C, *p*<0.001, n=13, unpaired *t*-test, Fig. 5I, left bars). The same comparison for warm spraying showed no statistically significant differences (Fig. 5I, right bars).

Most TRPM8-expressing free nerve terminals are located in the *stratum spinosum* and *stratum granulosum* of the epidermis (26). However, to understand whether and how subcutaneous (s.c.) temperature is affected by cold spraying, miniature temperature transponders were implanted s.c. above the thoracic spine in another group of mice. Albeit cold spraying significantly reduced dorsal fur temperature (FLIR measurements), dorsal s.c. temperature was much less affected (Fig. 5C, inset).

### Chickens respond to C-1 with a WDS-like behavior

The study of TRPM8-dependent WDS-like behaviors in birds has been limited by the insensitivity of avian TRPM8 to icilin. After identifying C-1 as a robust agonist for both recombinant and native cTRPM8 *in vitro* and demonstrating that rodents respond with vigorous WDS to C-1, we hypothesized that C-1 would elicit a putative TRPM8-dependent shaking behavior in chickens. Pharmacological preparations were s.c. (axillary) administered to chickens, as it was difficult to inject solutions i.p. while avoiding the abdominal air sacs.

C-1 (33 mg/kg) immediately evoked a range of behaviors: ample feather ruffling, preceding any vigorous body shaking, head and neck shaking, and finally, crowding in corners, often in a sleep-like behavior (Fig. 6A, i-iv, Fig. 6B, Movie S5 and S6). Head and neck shaking, with a maximum twist amplitude of about 180°was performed with closed eyelids (Fig. 6A, i). Feather ruffling was most visible around the neck, chest, and leg feathers, and could be seen from above by the large increase in feather volume (Fig. 6A, ii). A few seconds after feathers were ruffled, a vigorous shaking of the body occurred, with the wings slightly spread and flapping (Fig. 6A, iii). When the previously described behaviors ceased, the birds started pressing their chest against a corner or enclosure wall while having fluffed feathers, as if to protect themselves from cold (Fig. 6A, iv, Movie S6), and then entered a sleep-like state. This corner-crowding behavior occurred no later than 130 s after the injection and continued throughout the 15-min recording period. Two chickens were recorded for up to 30 min, a time they spent in the same sleep-like state in a corner.

**Fig. 6.**
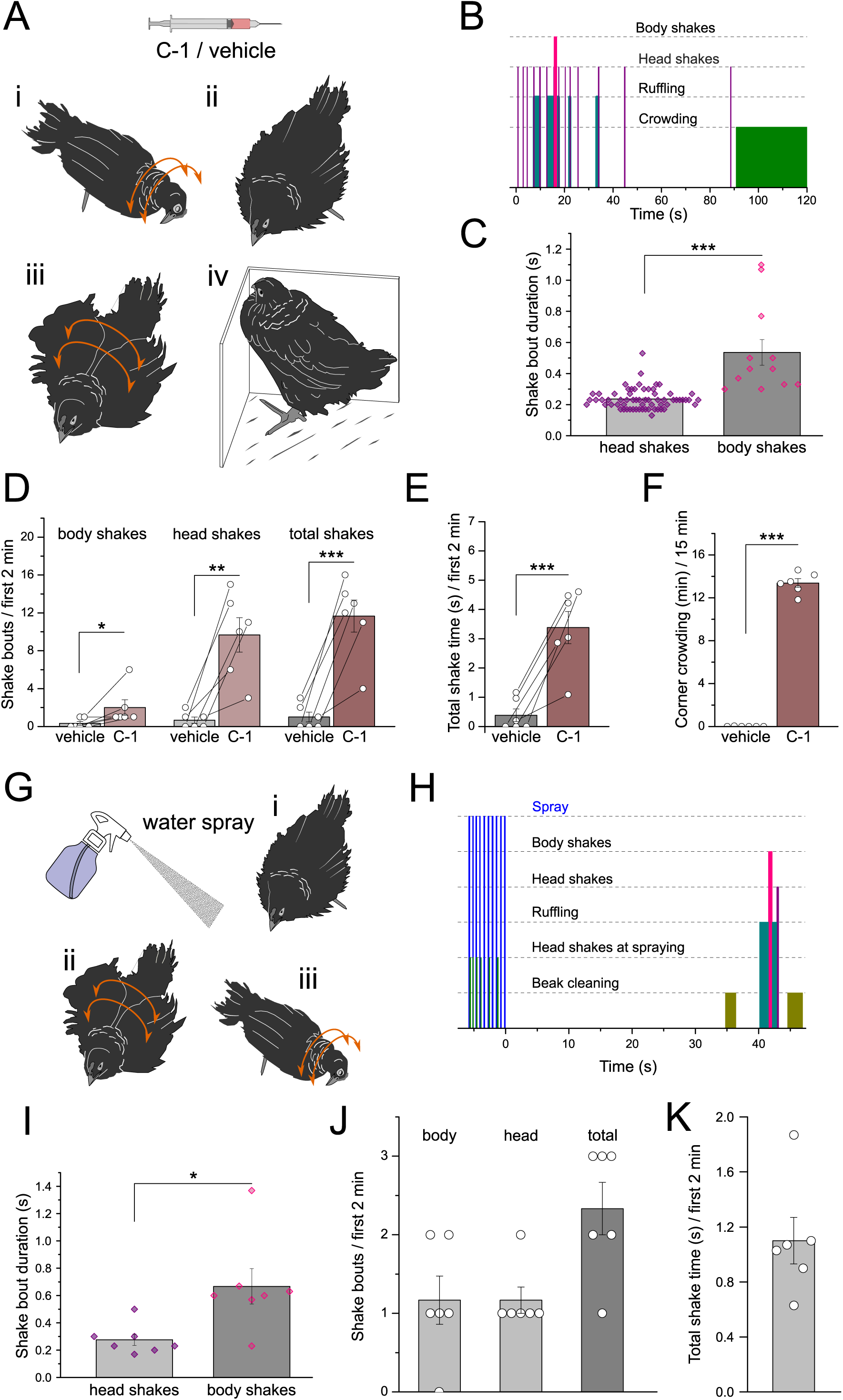
C-1 and water spraying evoke similar shaking behaviors in chickens. (A) Four typical behaviors recorded in juvenile chickens after s.c. C-1 administration (33 mg/kg). (**i**) Head shaking occurred with eyelids closed. (**ii**) Feather ruffling preceded all strong shakes. (**iii**) Vigorous body shakes were accompanied by moderate wing spreading and flapping (examples shown in Movie S5). (**iv**) Corner-crowding in a sleep-like state while pressing the chest against the walls of the arena typically occurred within 2 min after C-1 administration (examples shown in Movie S6). (B) A 2 min ethogram starting immediately after the C-1 injection. Typically, the body shakes were preceded by feather ruffling. (C) Body shake bouts elicited by C-1 were significantly longer than head shake bouts (***, *p*<0.001, *t*-test, unpaired, 64 head shakes and 12 body shakes, pooled from 6 chickens). (D) The number of shake bouts counted over the first 2 min after injection. Body, head, and total number of shake bouts were significantly higher in chickens injected with C-1 compared with the same chickens receiving only vehicle (***, *p*<0.001; **, *p*<0.01; *, *p*<0.05, *t*-test, paired, n=6 chickens). (E) The total shake time in the first 2 min after the injection (the same experimental set as in D), C-1 compared to vehicle (***, *p*<0.001, *t*-test, paired, n=6 chickens). (F) Total time spent during corner-crowding, pressing the chest against the walls of the arena. This behavior characterized every C-1 injected chicken, while it was absent in vehicle-injected chickens (***, *p*<0.001, *t*-test, paired, n=6 chickens). (G) All behaviors described above (in A), except for corner-crowding, were present in laboratory water-sprayed chickens (Movie S9), as well as in farmhouse chickens splashed with water (Movie S7). (H) A short ethogram showing the behaviors of a chicken sprayed with water at room temperature (∼25 °C, 10 sprays in quick succession). (I) In water-sprayed chickens, body shake bouts were significantly longer than head shake bouts (*, *p*<0.05, *t*-test, unpaired, 7 head shakes and 7 body shakes, pooled from 6 chickens). (J) The number of shake bouts (body, head, and total) in the first 2 min from spraying (n=6 chickens). (K) The total shake time in the first 2 min after water-spraying (for the same experimental set as in J, n=6 chickens). **Figure 6 – Source data 1:** shake bout duration, number of shake bouts, total shake time and corner crowding behavior duration for Figure 6C-F.

Comparing head with body shakes in the 6 chickens treated with C-1, the elicited head shake bouts were more frequent and significantly shorter than body shake bouts (Fig. 6C).

C-1 elicited significantly more body, head, and total shake bouts than the vehicle, and a longer total shaking duration in the first 2 min, before chickens entered the sleep-like state (Fig. 6D, E). The few shaking movements induced by the vehicle can probably be explained by the temperature of the injected volume, which could not be precisely controlled and was therefore lower than the axillary s.c. temperature of the chickens. Corner-crowding and the sleep-like state were not present in vehicle-treated chickens, in contrast to the long duration of these behaviors in the same chickens receiving C-1 (Fig. 6F).

### Chickens respond to water-spraying with a WDS-like behavior

We first examined the shaking behavior of domestic chickens by splashing adult farm chickens with water or sprinkling them with dust and found that the birds exhibited robust behaviors starting with feather ruffling, followed by vigorous body shaking and ending with neck twisting (Movie S7, S8). Following these observations, to closely investigate shaking in response to naturalistic stimuli, we water-sprayed the chickens under controlled laboratory conditions.

Each chicken was sprayed 10 times in rapid succession (∼0.9 ml per actuation) with water at room temperature (∼25 °C). Chickens reacted during spraying with very short startle-like head shakes, which were not included in further analyses. Feather ruffling, body and head shaking, and an additional behavior, beak cleaning on the floor, followed soon after spraying (Fig. 6G, H, Movie S9). The corner-crowding in the sleep-like state observed in C-1 injected chickens was absent in the spray experiments. Similarly to the response to s.c. C-1, the average duration of the body shake bouts was significantly longer than for head shake bouts (Fig. 6I). The total number of shake bouts in the first 2 min after water-spraying, and their total time, had lower values than for the equivalent movements elicited by C-1 injections (Fig. 6J, K). Altogether, the above suggest that feather wetting-evoked shaking behaviors in chicken can be reproduced with the TRPM8 agonist C-1.

## DISCUSSION

Here, we have compared the chemically induced shaking behavior in mammals and birds with that induced by natural water-spray stimuli. We were able to show that chemical stimulation of TRPM8 elicits WDS-like behaviors in all tested species, and that TRPM8 contributes to the cold water evoked WDS in mice.

### A TRPM8 agonist that can elicit shaking

Since the cloning of the cold and menthol receptor TRPM8 (25), a variety of synthetic small molecules have been validated as agonists of the channel, often with higher potencies than menthol (27). However, of this large number of potent agonists, until recently only icilin was known to induce robust WDS in rodents. Here we show that C-1, a DIPA agonist of TRPM8, is capable of eliciting robust WDS-like responses in awake rodents and birds.

We demonstrated that C-1 is a potent and efficacious agonist of the icilin-insensitive chicken TRPM8, as well as TRPM8 orthologues from mammals (Fig. 1B). The menthol-insensitive Y745H hTRPM8 mutant was essentially insensitive to C-1 (Fig. 1C), confirming that the new agonist binds the same tyrosine residue as most TRPM8 agonists (17, 18, 28). Therefore, icilin-like or Y745-independent TRPM8 binding is not required for the WDS-inducing properties of C-1.

The concentration-response dependence showed a higher sensitivity of cTRPM8 to C-1, compared to r/hTRPM8. Previously, cTRPM8 was shown to be more sensitive to menthol than mouse TRPM8 (19, 20). We now show that chicken TRPM8 is much more sensitive to C-1 than to the menthol derivative WS-12, whereas in mammalian orthologs, the order of sensitivity is reversed (Fig. 1H, Fig. S1).

AMTB abolished both calcium transients and whole-cell currents following activation of recombinant TRPM8 by C-1. Chicken TRPM8 recovered more rapidly after inhibition than rat and human TRPM8, which were more sustainably inhibited by AMTB (Fig. 1E, Fig. S1B).

Whole-cell current recordings in cells transfected with each of the three TRPM8 orthologs confirmed the pharmacological properties shown with calcium imaging.

Simultaneous stimulation of recombinant TRPM8 with decreasing temperature ramps and agonists revealed a diffusion-limited dependence of TRPM8 open state conduction once the maximum open probability of the channel was reached and its temperature-dependent gating became inconsequential (Fig. 1H, Fig. S1E). This property allows testing agonist efficacy: the higher the temperature at which the TRPM8 current starts to decrease during a cooling ramp, the more efficacious the agonist is. Our results suggest that 10 µM C-1 can maximally activate cTRPM8, while 10 µM WS-12 can do the same for r/hTRPM8. A variety of ion channels have similarly low Q_10_ values for open channel conductance (29) as those computed here for the current amplitudes of TRPM8 orthologs stimulated by C-1 or WS-12 (∼1.3).

In experiments performed on cultured DRG neurons, we showed that their sensitivities to WS-12 and C-1 overlap to a large extent in all tested species. In rat DRG cultures, our results indicate that both 10 µM C-1 and 5 µM WS-12 failed to recruit all TRPM8-expressing rat neurons, whereas 100 µM C-1 likely stimulated all neurons (Fig. 2A). The overlap between the sensitivities to WS-12 and C-1 was lower in experiments done in DRG cultures from WT mice probed with Fura-2 imaging (Fig. 2C iv). Interestingly, the absence of TRPM8 revealed a substantial population of neurons displaying calcium transients in response to WS-12 (5 µM). Although WS-12 has been generally considered a selective TRPM8 agonist, this was based on recordings from *Xenopus* oocytes expressing recombinant TRP ion channels (30). Our current results, questioning the TRPM8 selectivity of WS-12 in mouse DRG neurons, are in agreement with our previous findings (31). It can therefore be concluded that sensitivity to C-1 is a more reliable indicator of functional expression of TRPM8 than the relatively nonselective effect of WS-12 in mouse DRG neurons.

The WDS induced by icilin depend on the peripheral nervous system, as injections into the lateral ventricles or striatum of rats did not elicit WDS (4), whereas injections into the cisterna magna elicited head shaking only, with a dose dependence shifted to higher doses compared to i.p. administration (2). Furthermore, TRPM8 is expressed to a much lesser extent in the brain than in the peripheral nervous system (32).

In our experiments, both icilin and C-1 elicited robust WDS after i.p. injections in rats, the preferred animal model for studying WDS (Fig. 3A-D) and were reported to respond to icilin with more frequent WDS bouts than C57BL/6 mice (9, 13). Our results in rats are consistent with a previous report of DIPA-induced WDS in anesthetized rats (16). We have also tested in rats another potent agonist, WS-12 (30, 33), without any success in inducing WDS. The uncommon property of TRPM8 agonists of inducing WDS is likely due to a favorable trade-off between potency and *in vivo* bioavailability, which allows quick access to cutaneous nerve terminals expressing TRPM8 at sufficiently high concentrations. Therefore, the solubility of these compounds, characterized by logP or logD at physiological pH, should also be considered when searching for suitable WDS-inducing TRPM8 agonists. High doses of menthol were reported to induce WDS in rats, albeit of a shorter duration than icilin, while also inducing writhing behavior (a sign of pain), ataxia, and loss of righting reflex (23). Another synthetic TRPM8 agonist reported to cause sporadic WDS in rats is M8-Ag (34). The predicted logD (at pH 7.4) is 2.23 for icilin, 2.66 for C-1, 2.79 for M8-Ag, 3.14 for menthol, while for WS-12 it is 5.02 (as predicted by ACD/PhysChem Suite), suggesting that a logD in the range of 2 to 3 is important for eliciting WDS after i.p. administration.

We also triggered WDS by spraying rats with water at room temperature. These WDS bouts were less frequent than the chemically-elicited ones (Fig. 3).

In mice, we demonstrate the TRPM8 dependence of WDS triggered by C-1 using *Trpm8-/-* mice that did not shake in response to i.p. administered C-1. This is consistent with previous findings of icilin-induced WDS and jumping in mice (9, 10). Besides WDS, another striking difference between WT and *Trpm8-/-* mice, when injected with C-1, was the grooming duration, which was significantly longer in WT mice, with a more frequent crouched posture in these mice (Fig. 4).

### The role of TRPM8 in the fast shaking behavior of water-sprayed mice

We felt it was necessary to approach the study of TRPM8-dependent shaking in a more naturalistic manner, by looking at how TRPM8 function affects the shaking of wet terrestrial mammals. Considering the importance of TRPM8 in cold sensing, we investigated the role of temperature in inducing WDS by spraying WT and *Trpm8-/-* mice with water at two temperatures. Most importantly, we found that the latency of the first WDS after spraying was significantly shortened in a cooling-dependent manner in WT mice only, indicating that TRPM8 controls the urgency of shaking in mice (Fig. 5G, H). We also found that in the first 30 s and 1 min after spraying, WT mice shook more when sprayed with cold than with warm water, whereas this was not the case for the *Trpm8-/-* mice (Fig. 5E, F).

This demonstrates the importance of a dry coat to avoid heat loss that is likely to occur upon contact with cold water. TRPM8 activation likely acts as a warning signal of potentially critical body heat loss due to wetness and reduces the latency of the first shake compared to mechanical stimulation alone (Fig. 7). In contrast to mechanosensitivity, which does not provide information about heat loss due to wetness, temperature sensitivity via TRPM8 helps to detect the temperature drop of the skin, which depends on both the water temperature and its mass.

**Fig. 7.**
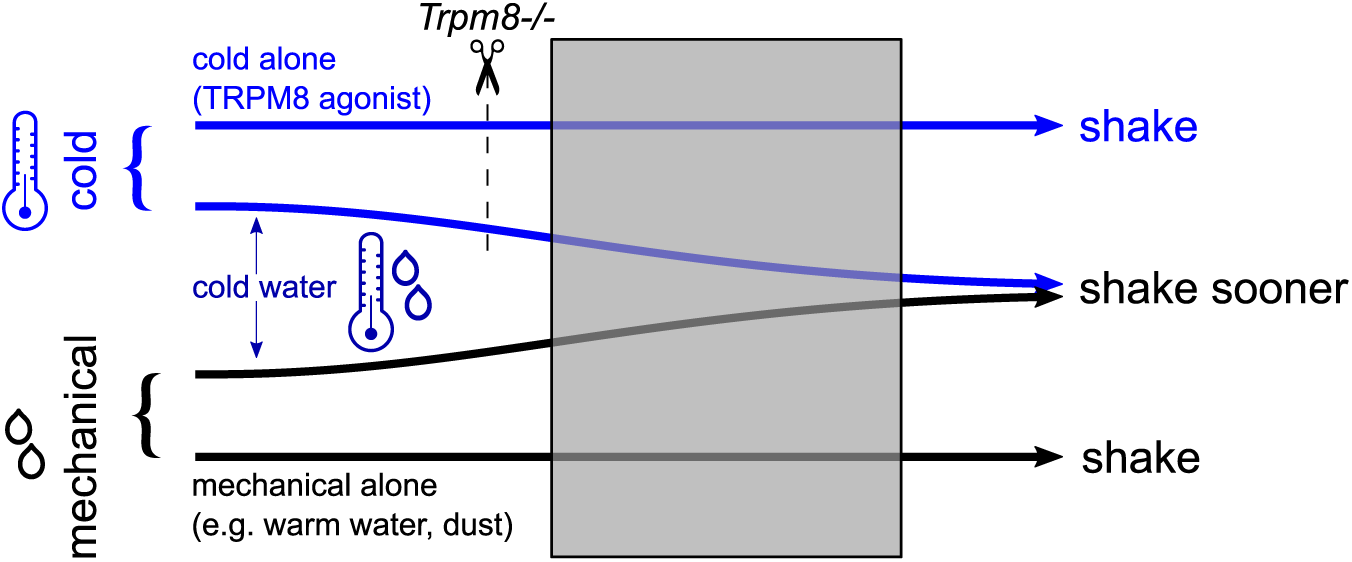
Proposed model for the TRPM8-dependency of the shaking behavior. Potent and bioavailable agonists of TRPM8, such as C-1, can elicit shaking by themselves, in the absence of any mechanical stimulation (top horizontal arrow). This was shown in the current study in both rodents and chickens. Mechanical stimulation alone (e.g., warm water, dust) can evoke shaking in the absence of cold stimulation (bottom horizontal arrow). Combining the two stimuli, as in cold water spray (middle horizontal arrows), modulates the shaking response by decreasing the latency of the first shake compared to mechanical stimulation alone. TRPM8 absence (scissors symbol) abolishes the shaking elicited by TRPM8 agonists and the latency reduction in cold-evoked shaking.

Paradoxically, in the interval between the first and second minute, when the dorsal fur temperature difference between cold and warm spraying was much reduced, *Trpm8-/-* mice shook more after cold than warm spraying (Fig. 5D, E). This could indicate that C57BL/6J-based *Trpm8-/-* mice possess a TRPM8-independent and delayed cold-sensing mechanism. Further experiments are needed to elucidate the identity of the cold– and mechano-sensitive receptors involved in TRPM8-independent shaking responses to wet fur.

### TRPM8 involvement in avian shaking behaviors

Bird shaking behaviors are trivially known but their mechanisms have hardly been studied (7). Here we report the elicitation of WDS-like behavior in birds by pharmacological agents. The qualitative inspection of the elicited shaking behavior revealed slightly different behaviors between chickens and rodents, as in chickens the head and body shaking appeared to be distinct and not necessarily part of the same sequence (Fig. 6B, Movie S9). In chickens, the duration of individual shake bouts was longer for body shakes than for head shakes. This feature was common to both pharmacologically (Fig. 6C) and spray-evoked shaking (Fig. 6I).

Body shaking was facilitated by slight movements of the wings. The gradual propagation of the twisting movement from the head to the tail, as seen in mammals, was not as evident in chickens (Movie S9), owing perhaps to vertebral fusion, common in birds. In chickens, vertebral fusion is age-dependent and occurs at several levels, particularly the thoracic (forming the notarium) and thoracic to caudal levels (forming the synsacrum) (35). Our experiments were conducted with young birds, which most likely still had free vertebrae, but the same shaking behavior is also observed in older chickens (Movie S7). It is noteworthy that body shaking is a much less frequent event in chickens compared to rats or mice: the number of body shake bouts was 1-2 in most birds tested, regardless of stimulation type (C-1 or water-spraying, Fig 6B, H).

Interestingly, C-1 injections also elicited feather ruffling and, after about two minutes, a long-lasting crowding against the enclosure corners and a sleep-like state with fluffed feathers. These are known heat-loss limiting behaviors encountered in chickens during cold exposure (36, 37). This is consistent with a prolonged *in vivo* effect of C-1 on chicken TRPM8.

The same corner-crowding and sleep-like behaviors did not occur in chickens sprayed with water at room temperature. This suggests that the stimulation did not decrease body temperature enough to evoke behaviors associated with cold weather exposure (36, 37). This is also supported by the lower number of shake bouts and shorter total shake time in sprayed chickens, compared to chickens injected with C-1 (Fig. 6D, E, J, K).

### The evolution of shaking and its relation to TRPM8

How shaking behaviors and their recruitment of TRPM8 evolved can only be speculated upon, but it is nevertheless a fascinating question involving the evolution of hairs and feathers, spine twisting capacity, and TRPM8 itself.

Having shown that mammals and birds share a TRPM8-dependent shaking behavior, the question naturally arises as to how this behavior evolved – is it conserved, or did it evolve at least twice? The most recent common ancestor of mammals and birds – an early amniote – had no capacity for axial twisting of the thoracic spine (38, 39) and probably no need to shake, as it likely had a self-cleaning hydrophobic integument, as some modern lizards (40, 41). On the other hand, it likely already took advantage of TRPM8’s sensitivity to cold, which was acquired as the first tetrapods emerged, long before hairs and feathers evolved (42).

The convergent evolution of a strikingly similar shaking behavior in furred and feathered animals reveals the importance of keeping body coverings dry and clean. Shaking around the spinal axis was an efficient way to achieve this.

While flying birds may also benefit from removing water from their feathers to become lighter (even during flight, (7)), all birds appear to have similar shaking behaviors. Skeletal and muscular adaptations for flight are not a prerequisite for shaking in birds, as Palaeognathae (ratites) can shake to remove water from their plumage (emu: (43, 44); cassowary: (45)), despite a nearly absent furcula, non-keeled sternum, and reduced pectoral girdle musculature (46).

It is reasonable to infer that, if shaking and grooming were not permitted by the skeleton, the survival of an extensively furred or feathered, non-aquatic endothermic tetrapod would require high energy costs for thermoregulation after cold water soaking of its coat, especially in small animals (47).

Since proper hygroreceptors have only been found in invertebrates (48), wetness sensing in furred or feathered animals is thought to be synthetic, arising from mechanical and cold sensing, by analogy with human perception (49). It is known that mechanical stimulation alone (e.g., coat touching, dust pressure following dust bathing) evoke WDS in mammals and birds (50), see examples, Movie S3, S8).

To our knowledge, there is yet no evidence that natural cold stimuli (i.e., exposure to cold air) can trigger WDS by themselves. One explanation may lie in the adaptation to low ambient temperatures, facilitated by the comparatively slow variation of air temperature, in contrast to raindrops and bathing, which produce a much faster surface temperature change. Consequently, it is unlikely that the animals in their natural environment are confronted with circumstances in which the temperature decreases rapidly without a mechanical stimulus.

Regardless of the conceivable dissociation of strong cold and mechanical sensing, we believe that shaking in response to wet-cold conditions is an important thermoregulatory behavior in mammals and birds. Our research shows that the cold sensor TRPM8 controls the urgency to shake, providing a rapid regulatory response for avoiding excessive heat-loss.

## MATERIALS AND METHODS

### Animal models and behavior

#### Rats

The animals used were 17 Wistar rats (purchased from the Nicolae Simionescu Institute of Cellular Biology and Pathology), 4-8 weeks old at the time of experiments. The weight of rat males was 193.7 ± 18.3 g (mean ± SEM, n=9), and that of females 178.0 ± 11.8 g (mean ± SEM, n=8). The rats were housed in cages of 2 animals and kept on a 12-h light-dark cycle with the experiments being conducted during the light phase. Rats were supplied with water and food *ad libitum*.

### *Trpm8^-/-^* mice group

The animals used were wild-type C567BL/6 (purchased from Jackson Laboratory, https://www.jax.org/strain/000664), called WT, and TRPM8 knock-out mice (*Trpm8^-/-^* mice on C56BL/6 background, (51); also from Jackson Laboratory, https://www.jax.org/strain/008198), called further *Trpm8-/-*, in total 20 mice of both sexes, aged 12-22 weeks (118.2 ± 7.0 days, n=20). The weight (mean ± SEM) of the WT mice was 27.8 ± 1.2 g (males, n=6) and 22.0 ± 1.0 g (females, n=4), while that of *Trpm8-/-* mice was 27.2 ± 0.5 g (males, n=6) and 19.8 ± 0.5 g (females, n=4). Mice were housed up to 5 per cage, kept on a 12-h light–dark cycle, and were supplied with water and food *ad libitum*.

### Trpm8^EGFPf/EGFPf^ mice group

The animals used were wild-type (WT) C57BL/6J (purchased from Jackson Laboratory, RRID: IMSR_JAX:008198) and TRPM8 KO mice (*Trpm8^EGFPf/EGFPf^* on C57BL/6 background, (9); a generous gift from Prof. Ardem Patapoutian, The Scripps Research Institute), called further *Trpm8-/-*, in total 26 mice of both sexes, 8-16 weeks old (83.0 ± 2.4 days, mean ± SEM, n=26). The weight (mean ± SEM) of the WT mice was 25.6 ± 0.7 g (males, n=7) and 22.4 ± 0.5 g (females, n=6), while that of *Trpm8-/-* mice was 25.1 ± 0.6 g (males, n=7) and 22.0 ± 0.8 g (females, n=6). Mice were housed in group cages inside a ventilated cabinet (Scantainer, Scanbur Technology A/S, Karlslunde, Denmark), kept at constant temperature (22 °C) and humidity (55 %) on a 12-h light-dark cycle and were supplied with water and food *ad libitum*.

#### Chickens

The animals used were Plymouth Rock barred chickens (purchased from Agromar Balotesti) of both sexes, 6 animals in total (estimated as 3 males and 3 females), all of the same age, 30-38 days of age during the study. Chickens were housed together in a large bird cage, at about 23 °C and 40% relative humidity, on a 12-h light-dark cycle and were supplied with water and food *ad libitum*. The sex of the chickens was estimated based on head plumage color (52) and later by the development of the comb and wattles in males. The average mass of the chickens during the C-1 experiments was 644.2 ± 40.1 g (mean ± SEM, n=6).

#### Behavioral experiments in rats

The procedures for investigating the effects of i.p. C-1 and water spraying in rats were approved by the Research Ethics Committee of the University of Bucharest (No. 75/20.04.2019) and complied with EU Directive 2010/63/EU and the Romanian law 43/11.04.2014 on laboratory animal protection. The recording arenas consisted of bottomless transparent acrylic boxes (length=21 cm; depth=16 cm; height=31 cm) with polycarbonate plastic floors. All rats in the spraying experiments were habituated with the enclosures during 3 sessions of 15 min each over 2 days before experiments. Icilin (1 mg/kg, in 100 µl solution/rat) was administered i.p. dissolved in polyethylene glycol 400 (PEG, Kollisolv 06855, Sigma-Aldrich), while C-1 (33 mg/kg, in 100 µl solution/rat) was dissolved in Ringer solution. This contained (in mM): NaCl, 140; KCl, 4; CaCl_2_, 2; MgCl_2_, 1; HEPES, 10; NaOH, 4.54 (pH 7.4 at 25 °C adjusted with NaOH). In control experiments, the animals received only the vehicle of each active compound. WS-12 (up to 33 mg/kg) was dissolved in either PEG or mixtures of two or three solvents (from ethanol, DMSO, PEG, Tween 20 and sunflower oil). The boxes and floors were completely dry and clean before each experiment. The room temperature in the icilin and C-1 experiments was kept between 24 and 26 °C. In the spraying experiments, the average room temperature and relative humidity were 24.0 ± 0.4 °C (mean ± SEM, n=3) and 36%, respectively. Each rat was sprayed with water at room temperature 20 times in quick succession, with 0.9 ml of water per spray, from a distance of ca. 36 cm. Each spray lasted less than 0.5 s and the next one was delivered immediately after. The total spraying duration was 8.1 ± 0.2 s (mean ± SEM, n=6), measured by analyzing the peak spectral density of the sound between the first and last spray, using the spectrogram feature in Solomon Coder (version: beta 19.08.02, RRID: SCR_016041).

Rat behavior was recorded using a color camera from the side or above the arenas, for 30 min after the injection, at a resolution up to 2224 x 1080 pixels at 25 or 30 frames per second (fps). After 30 min elapsed, the animals were returned to their home cages. At the end of the spraying experiments, the excess water was gently swept from the rat’s fur with paper towels.

#### Behavioral experiments in mice treated with C-1

The procedures for investigating the effects of i.p. C-1 on mice were approved by the UC Davis Institutional Animal Care and Use Committee (IACUC), Protocol for Animal Care and Use #22313. All animals were habituated (1 h/day) to the recording arenas (open cylindrical restrainers, diameter=12 cm; height=25 cm) for 3 days before testing. The enclosures sat on a plastic floor and during the recordings were covered with a transparent acrylic plate. The laboratory temperature was controlled and maintained in the range of 20-22 °C, while the relative humidity was in the 30-40% range. C-1 (33 mg/kg in 10 ml/kg of Ringer solution, contents described above) was injected i.p. In control experiments, the same animals received only the vehicle. The behavior was recorded from above the arenas for 30 min after the injection, at a resolution of 1920 x 1080 pixels, at 25 fps. After 30 min elapsed, the animals were returned to their home cages.

#### Behavioral experiments in water-sprayed mice

The procedures investigating the behavioral responses in mice following warm and cold water spraying, were performed according to regulations of animal care and welfare and approved by the Animal Protection Authority of the District Government of Lower Franconia (Würzburg, Germany) and the ethics committee of Friedrich-Alexander University Erlangen-Nürnberg under the file number 55.2.2-2532-2-1014. Mice were habituated in a bottomless transparent polycarbonate box (length=15 cm; depth=14 cm; height=20 cm) during three sessions of 15 min each, over 2-3 days. The box sat on a polypropylene floor. The box and the floor were completely dry and clean before each experiment. On the day of the experiment, the mice were left in their home cages in the room where the experiments were performed, for at least 20 minutes, to acclimatize to room conditions. The average room temperature and relative humidity during the experiments were 24.8 ± 0.1 °C and 40.2 ± 0.5%, respectively (mean ± SEM, n=52 experiments).

The mice were sprayed with a pump bottle with the nozzle adjusted to deliver ∼0.85 ml per each actuation. Each spray lasted less than 0.5 s and the next one was delivered as soon as possible. Each mouse was sprayed 4 times in quick succession. The spraying duration was 1.61 ± 0.03 s (mean ± SEM, n=52 experiments), measured by analyzing the peak spectral density of the sound between the first and last spray, using the spectrogram feature in Solomon Coder. The warm and cold spraying duration averages were comparable: 1.57 ± 0.04 s and 1.65 ± 0.05 s, respectively (mean ± SEM, n=26 experiments).

Spraying targeted the rostral back area. On the ground (from the height of ca. 25 cm), the shape of the sprayed area was an oval with extreme diameters of ca. 18 and 11 cm. Each mouse was sprayed with water at two temperatures, measured inside the bottle with a thermocouple, and on the surface of the mouse, using a thermal imaging camera (T420bx, FLIR Systems Inc, FLIR(R) Systems GmbH, Frankfurt, Germany). Distilled water was warmed and cooled with two liquid baths programmed at 62 and 0.5 °C, respectively, then transferred in the pump bottle, which was equipped with a thermocouple wire connected to a digital thermometer. At 58 °C in the bottle, the measured fur temperature on the back of the mouse was in the range of 27 to 38 °C. At 4 °C in the bottle, the fur temperature was in the range of 11 to 20 °C.

Each mouse was sprayed inside the box where it was video-recorded. The recordings started with spraying and ended 15 min afterwards. Two cameras were used simultaneously: a high-definition color camera recording through the front panel of the box, and the FLIR camera, recording from above. The front camera was used to record video files of 2560 x 1440 pixels at 30 fps, while the FLIR camera was used to record radiometrically at 320 x 240 pixels and 30 fps. The thermal infrared recording was analyzed with the FLIR Tools Professional and FLIR Thermal Studio programs (Teledyne FLIR LLC, Wilsonville, OR, USA, RRID:SCR_016330) to measure the average fur temperature on the back of each mouse, using the polygon selection tool available in FLIR Thermal Studio.

Each mouse underwent a single spray experiment per day and most mice were sprayed during two consecutive days. The order of exposure to warm and cold spray experiments was alternated in all animal groups – so that approximately half of the mice were sprayed first with warm water. If the temperature measured immediately after the end of last spray was not in the interval desired for neutral-warm (27 – 37 °C) or cold (12 – 19 °C) temperatures, the experiment was repeated the following day. The average number of spray trials to which a mouse was exposed was 1.23 ± 0.10 (minimum 1, maximum 3) for warm spray and 1.38 ± 0.13 (minimum 1, maximum 3) for cold spray (mean ± SEM, n=26). The average temperature difference between warm and cold spray per mouse was 17.4 ± 0.5 °C (mean ± SEM, n=26). After 15 min elapsed, the mouse was removed from the box, the excess water from its fur was gently swept with a paper towel, and the mouse was returned to its home cage.

A different group of mice, 3 WT and 3 *Trpm8-/-*, all males, 13-14 weeks old (95.3 ±1.0 days, mean ± SEM, n=6), weighing 25.8 ± 0.9 g, was implanted s.c. with miniature transponders (temperature microchip UCT-2112, Unified Information Devices, Lake Villa, IL, USA) under the back skin, above the middle thoracic spine, using the manufacturer supplied microchip injector and hypodermic needles. The measured s.c. temperature was read with a receptor unit (URH 1HP reader, Unified Information Devices) placed under the floor of the enclosure and connected to a computer. Temperature acquisition started at least two minutes before cold spraying. The interval between consecutive data points was typically one second but was dependent on the position of the mouse inside the enclosure, relative to the receptor. One mouse was excluded from analysis due to a technical problem that introduced a long delay (over one minute) in acquiring the temperature after spraying.

#### Behavioral experiments in chickens

The procedures for investigating the effects of s.c. C-1 and water-spraying in chickens were approved by the Research Ethics Committee of the University of Bucharest (No. 75/20.04.2019) and complied with the EU Directive 2010/63/EU and the Romanian law 43/11.04.2014 on laboratory animal protection. Chickens were habituated to the arena, a bottomless transparent polycarbonate box (length = depth = height = 30 cm), during 2 sessions of 15 min each, over 2 days. The floor of the box was made from polycarbonate plastic covered with paper. The box and the floor were completely dry and clean before each experiment. Glass panels were used for covering the boxes during recordings, to discourage escape attempts.

In the spraying experiments, the average room temperature and relative humidity were 25.2 ± 0.1 °C and 44.1 ± 1.1% (mean ± SEM, n=6), respectively. Each chicken was sprayed with water at room temperature 10 times in quick succession, with ∼0.9 ml of water per spray, from a distance of ca. 25-30 cm. Each spray lasted less than 0.5 s and the next one was delivered immediately after. The spraying interval was measured using the Solomon coder spectrogram (as described above) and by visually following the stream of droplets in the recorded videos. The mean spraying duration was 4.5 ± 0.3 s (mean ± SEM, n=6).

C-1 (33 mg/kg) was dissolved in Ringer solution (see above) in a total volume of 1 ml/kg and s.c. injected under an axillary fold of skin. In control experiments the same animals received only the vehicle. The average room temperature in the C-1 experiments was 25.1 ± 0.3 (mean ± SEM, n=6). The behavior was recorded using a color camera from above the arena for up to 30 minutes after the injection, at a resolution of 2560 x 1440 pixels, at 30 fps. After the recordings, the chickens were returned to their home cages. At the end of the spraying experiments, the excess water on the plumage was gently swept with paper towels.

### Cell culture

#### Rat DRG

Rat DRG neurons were obtained from all spinal levels of adult male Wistar rats (purchased from the Nicolae Simionescu Institute of Cellular Biology and Pathology, Bucharest, Romania). Male rats (150–200 g; 8–10 weeks, n=4) were killed in a gradually increasing CO_2_ concentration followed by decapitation. The procedures were approved by the Research Ethics Committee of the University of Bucharest and complied with EU Directive 2010/63/EU and the Romanian law 43/11.04.2014 on laboratory animal protection. Upon excision from all spinal levels, DRGs were incubated with a mixture of 1.5 mg/ml collagenase type XI (C 7657, Sigma-Aldrich) and 3 mg/ml dispase (Gibco 17105-041, Thermo Fisher Scientific, Waltham, Massachusetts, USA) in the IncMix solution for 1 h at 37 °C and 5% CO_2_. Neurons were dissociated by gentle trituration with fire-polished Pasteur pipettes, plated on poly-D-lysine treated (0.1 mg/ml for 30 min) 25-mm borosilicate coverslips (0.17 mm thick) and maintained at 37 °C with 5% CO_2_ in a 1:1 mixture of DMEM and Ham’s F-12 medium (D 8900, Sigma-Aldrich), supplemented with 50 ng/ml mouse Nerve Growth Factor-7S (NGF, N 0513, Sigma-Aldrich), 10% horse serum and 50 µg/ml gentamicin. The cultures were used for experiments within 36 h after plating.

#### Mouse DRG

Mouse DRG neurons were excised from all spinal levels of adult male C57BL/6 mice (8 weeks, n=2, in-house breeding colony; originally from Jackson Laboratory, Bar Harbor, ME, USA, RRID: IMSR_JAX:000664), and *Trpm8^EGFPf/EGFPf^*mice (8 weeks, n=2) (9) a generous gift from Prof. Ardem Patapoutian, The Scripps Research Institute). All animal procedures were performed according to regulations of animal care and welfare (EU Directive 2010/63/EU) and approved by the Animal Protection Authority of the District Government (Ansbach, Germany). Mice were euthanized by breathing an increasing CO_2_ concentration. DRG were dissected out, transferred to DMEM containing 50 µg/ml gentamicin, treated with 1 mg/ml collagenase type XI (C 7657, Sigma-Aldrich) and 0.1 mg/ml protease (P5147, Sigma-Aldrich) for 30 min, mechanically dissociated with a fire-polished silicone-coated Pasteur pipette, and plated on poly-D-lysine treated (0.2 mg/ml) 10 mm diameter glass coverslips. DRG neurons were cultured in serum-free TNB 100 cell culture medium (F8023 Biochrom, Berlin, Germany) supplemented with TNB 100 lipid-protein complex (F8820, Biochrom) and streptomycin/penicillin (100 µg/ml). Mouse NGF 2.5S (N-240, Alomone Labs, Tel Aviv, Israel) was added at 100 ng/ml and the cells were kept at 37 °C and 5% CO_2_. Cells were used for experiments within 24 h.

#### Chicken DRG

For obtaining chicken DRG cultures, the procedures closely followed those used for culturing rat DRG neurons. Plymouth Rock barred chickens (purchased from Agromar Balotesti, Ilfov, Romania, 320-420 g; 4-5 weeks, n=3) were killed in a gradually increasing CO_2_ concentration followed by decapitation. The procedures were approved by the Research Ethics Committee of the University of Bucharest and complied with EU Directive 2010/63/EU and the Romanian law 43/11.04.2014 on laboratory animal protection. DRG neurons were obtained from all spinal levels. See above (rat sensory neuron culture) for the neuron dissociation and culturing procedures.

#### Heterologous expression of TRPM8 channels

Recombinant DNA was transiently transfected into HEK293T cells (RRID: CVCL_0063) using the jetPEI reagent (Polyplus-transfection S.A., Illkirch, France). Human TRPM8 (hTRPM8, RefSeq NM_024080.5), rat TRPM8 (rTRPM8, RefSeq NM_134371.3), and chicken TRPM8 (cTRPM8, GenBank OQ657222.1) were gifts from Dr. Gordon Reid, University College Cork (Ireland). Plasmids containing the hTRPM8 mutants Y745H were kind gifts from the laboratory of Dr. Viktorie Vlachova (Institute of Physiology, Academy of Sciences of the Czech Republic, Prague). All constructs were inserted in pcDNA3.1 vectors. Approximately 24 h after transfection, the cells were detached with trypsin-EDTA, washed with complete medium, centrifuged at 1000 g for 5 min, resuspended in medium and plated onto 25-mm borosilicate glass coverslips (0.17 mm thick), which had been treated with poly-D-lysine (0.1 mg/ml for 30 min, left to dry another 30 min). The culture medium was a 1:1 mixture of DMEM and Ham’s F-12 medium (D 8900, Sigma-Aldrich). For the concentration-response experiments, cells transfected with hTRPM8 or rTRPM8 were co-cultured at a 1:1 ratio with cells transfected with cTRPM8 to control for inter-dish variability in the calcium imaging experiments. The cultures were used for experiments within 36 h after plating.

### *In vitro* recordings

#### Single-wavelength calcium imaging

Intracellular non-ratiometric calcium imaging was used with HEK293T cells, and rat and chicken DRG neurons. Coverslips with attached cells were incubated for 30 min at 37 °C in ES containing 2 µM Calcium Green-1 AM (Invitrogen C3011MP, Thermo Fisher Scientific) and 0.02% Pluronic F-127 (Invitrogen P6867). Cells were washed and left to recover for 30 min before use. Coverslips were then mounted in a Teflon chamber (MSC TD, Harvard Apparatus, Holliston, MA, USA) on the stage of an Olympus IX70 inverted microscope and imaged with a 20× NA 0.75 objective. All solutions continuously superfused the cultured cells with a gravitational-driven 6-channel perfusion system through a miniature manifold (MM-6; Harvard Apparatus) at a rate of ∼0.75 ml/min. Temperature was controlled locally using a Peltier-based system (53). To select microscopic fields rich in cold-sensitive neurons, the cells were tested first with a mild cooling step. In order to measure the temperature experienced by the cells during the experiment, a miniature thermocouple (1T-1E, Physitemp, Clifton, NJ, USA) was placed very close to the imaged cells and the temperature was measured in real time with a digital thermometer (AZ 8851, TME, Łódź, Poland) connected to the imaging computer (Fig. 1A, lower part). Cells were illuminated with a 470 nm LED powered by a Dual OptoLED unit (Cairn Research, Faversham, UK) controlled by the Axon Imaging Workbench 2.2 software (AIW, Axon Instruments, Union City, CA, USA), which was also used for image acquisition and analysis. Fluorescence images were recorded with a CCD camera (Cohu 4910, Pieper GmbH, Schwerte, Germany) at a rate of 0.5-1 Hz and digitized to 8 bits. Regions of interest were used in AIW to mark individual cells. Background intensity was subtracted before computing the relative change of fluorescence (ΔF/F_0_). The average area under the ΔF/F_0_ curve (AUC), measured in seconds, during and before each stimulus, was used to discriminate cell responses. The threshold set to identify a cell response to chemical stimuli was set at the average baseline AUC plus its 2 standard deviations, for an interval equal to the ensuing stimulus duration. The difference values were rounded by truncation to 2 decimal places. To identify neurons responding to the cooling step, by excluding the small intrinsic Calcium Green-1 response to cooling (54), we set a threshold at 4 standard deviations from the center of the normal distribution fitting the small amplitude responses. Neurons with spontaneous activity, equal in amplitude with the activity recorded during chemical stimulation, were excluded from further analysis. Cells responding to 50 mM KCl were considered viable neurons. To analyze the data from the concentration-response experiments performed on coverslips with co-cultured HEK293T cells expressing either hTRPM8, or rTRPM8 and cTRPM8, icilin (10 µM) was applied at the end of experiments. The average amplitude (ΔF/F_0_) of the response of hTRPM8 and rTRPM8 was normalized to the response of cTRPM8 from the same culture dish. Unless otherwise stated, the experiments were performed at a constant temperature of 25 °C.

#### Ratiometric calcium imaging

Mouse DRG neuron cultures were loaded for 30 min at 37 °C with 3 µM Fura-2 AM (Invitrogen F1221) in ES containing also 0.02% Pluronic F-127 and left to recover for about 10 min in ES before recording. Coverslips were then mounted on the stage of an Olympus IX71 inverted microscope and imaged using a 10× objective. As imaging was done with a low magnification objective, temperature selection of microscopic fields rich in cold-sensitive neurons was not necessary. Cells were continuously superfused with ES using a software-controlled 7-channel gravity-driven common-outlet system (56). Fura-2 was excited at 340 and 380 nm using a Polychrome V monochromator (Till Photonics, Gräfelfing, Germany), with a full width half-maximum wavelength range of 10 nm. Fluorescence emission was long-passed at 495 nm and pairs of images were acquired at a rate of 0.5-1 Hz with an exposure time of 10 ms, using a 12 bit digital CCD camera and processed off-line (Imago Sensicam QE and TILLvisION software, Till Photonics, Gräfelfing, Germany). After background intensity was subtracted, the ratio between fluorescence emitted when Fura-2 was excited at 340 nm and at 380 nm (R = F_340_ /F_380_) was computed. Responsive cells were identified using the method described above (non-ratiometric calcium imaging). Neurons with spontaneous activity, equal in amplitude with the activity recorded during chemical stimulation, were excluded from further analysis. Cells responding to 1 µM capsaicin, or 50 mM KCl, were considered viable neurons.

#### Patch clamp electrophysiology

For patch-clamp recordings, HEK293T cells transiently transfected with TRPM8 orthologs and EGFP (DNA ratio 3:1), were plated on 25 mm borosilicate glass coverslips coated with poly-D-lysine and used within 24 h. Whole-cell patch clamp currents were recorded using a WPC-100 patch clamp amplifier (E.S.F. Electronic, Göttingen, Germany), filtered at 2 kHz and digitized at 5 kHz using an Axon Instruments DigiData 1322A interface driven by pCLAMP 8.2 (RRID:SCR_011323, Molecular Devices, Sunnyvale, CA, USA). Capacitive transients were compensated using the R-series, C-slow and C-fast adjustments of the amplifier. The extracellular solution was nominally calcium-free, the same as the solution for calcium imaging, except it contained no CaCl_2_ and was supplemented with 1 mM EGTA. The intracellular solution was K^+^-free with low Cl^-^ and contained (in mM): 134 Cs gluconate, 6 NaCl, 1 MgCl_2_, 2 Na_2_ATP, 5 EGTA, 10 HEPES; pH 7.22 at 25 °C adjusted with CsOH. Borosilicate capillaries with filament (GC-150F-10, Harvard Apparatus, Holliston, MA, USA) were pulled using a horizontal micropipette puller (P-1000, Sutter Instruments, Novato, CA, USA) and the tip polished for resistances of 2 – 4 MΩ. The reference electrode was placed in the same intracellular solution as the recording electrode and connected to the bath through an agar bridge. Currents were recorded either at –80 mV or during voltage ramps, from −100 mV to 100 mV (400 ms every 2 s, holding at −80 mV). During the experiments, the temperature inside the bath was controlled via a custom-made Peltier-driven perfusion system (53, 55) and cooling ramps were triggered using pCLAMP or a custom-written software. The temperature was measured as above (see calcium imaging) before and after the recording, at the place where the cell had been located. Unless otherwise stated, the recordings were performed with the temperature fixed at 25 °C. The representative current traces shown in the figures were decimated with a factor of 1000 in Clampfit.

#### Solutions and chemicals

The standard extracellular solution (ES) used in all experiments contained (in mM) NaCl, 140; KCl, 4; CaCl_2_, 2; MgCl_2_, 1; HEPES, 10; NaOH, 4.54; and glucose, 5 (pH 7.4 at 25 °C). The solution for DRG incubation (IncMix) contained (in mM): NaCl, 155; K_2_HPO_4_, 1.5; HEPES, 5.6; NaHEPES, 4.8; glucose, 5; and gentamicin 50 mg/ml. Drugs were added from the following stock solutions: (2S,5R)-2-isopropyl-N-(4-methoxyphenyl)-5-methylcyclohexanecarboximide (WS-12, Tocris #3040, Bio-Techne Ltd, Abingdon, UK) 5 mM in ethanol; 1-diisopropylphosphorylheptane (Cryosim-1, C-1, CAS Registry Number 1487170-15-9, supplied by E.T. Wei) 1, 10 and 100 mM in H_2_O; 1-(2-hydroxyphenyl)-4-(V3-nitrophenyl)-l,2,3,6-tetrahydropyrimidine-2-one (icilin, Tocris #1531) 10 mM in DMSO, 4-(3-chloro-pyridin-2-yl)-piperazine-1-carboxylic acid (4-tert-butyl-phenyl)-amide (BCTC, Tocris #3875) 10 mM in DMSO, N-(3-aminopropyl)-2-{[(3-methylphenyl)methyl]oxy}-N-(2-thienylmethyl)benzamide hydrochloride salt (AMTB, Tocris #3989) 10 mM in H_2_O, (1R, 2S, 5R) – (-) menthol (M278-0, Sigma-Aldrich, Saint Louis, MO, USA) 200 mM in ethanol, capsaicin (M2028, Sigma-Aldrich) 5 mM in ethanol, allyl isothiocyanate (AITC, #377430, Sigma Aldrich) 100 mM in DMSO. Fresh stock solutions of AITC and for all working drug dilutions were prepared on the day of the experiment. All unspecified chemicals were from Sigma-Aldrich.

### Statistical analysis

Data analysis and statistical tests were performed with either OriginPro 8 (RRID: SCR_014212, OriginLab Corporation, Northampton, MA, USA) or GraphPad Prism 9 (RRID: SCR_002798, Dotmatics, Boston, MS, USA). Data are presented as mean ± SEM. For illustration purposes, the averaged ΔF/F_0_ and F_340_/F_380_ were smoothed using a Savitzky-Golay filter set at a window length of 5-7. All data sets were found normally distributed according to the Kolmogorov-Smirnov normality test. Comparisons between two samples were performed using the two-tailed Student’s *t*-test (paired or unpaired, as indicated). Ion current densities elicited by one agonist at several temperatures in the same cells were compared using one-way repeated measures ANOVA. Current densities elicited by two agonists at two temperatures were compared with a repeated measures two-way ANOVA followed by Tukey’s post-hoc test. Responsiveness to TRPM8 agonists in DRG neurons from WT and *Trpm8-/-* mice was compared with the chi-square goodness-of-fit test for frequencies. Behavioral parameters measured at multiple time points were compared within or between groups with a repeated measures two-way ANOVA followed by Tukey’s post-hoc test, with the independent variables set as time and genotype, or time and spraying temperature.

Simple linear regression was used to estimate the relationship between the shaking latency and back fur temperature of water-sprayed mice for each genotype (WT and *Trpm8-/-*). A value of *p*<0.05 was considered statistically significant.

The dose-response curves for non-ratiometric imaging were fitted using the Hill equation (without 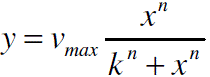 weighting), where *v*_max_ is equivalent to the maximal response amplitude, *k* to EC_50_ and *n* is the Hill coefficient. Data points that showed obvious desensitization (smaller response at 100 µM than at 10 µM C-1) were 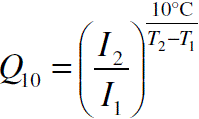 excluded from the fitting data sets. The Q_10_ coefficients, defined as were computed for linear regions of maximal slope spanning at least 2 °C when analyzing log(-current) against temperature.

Linear fitting was used to find the slope and Q_10_ was calculated as 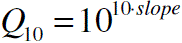 (57).

Predicted logP values were retrieved from ACD/Labs Percepta Platform (PhysChem Module) computed properties, available in the ChemSpider database.

The behavioral analysis was performed by investigators blind to all experimental conditions, using Solomon Coder or BORIS (RRID: SCR_021509, (58)). A shaking event in rats and mice was considered any visible head, neck, or body rotation. Shaking behavior was similarly scored in chickens, whereas, to avoid scoring spontaneous head twitches, we set a head shake bout duration threshold of 0.1 s.

Vector graphs were exported and compiled together with illustrations in InkScape 1.0 (RRID: SCR_014479).

## Acknowledgments

The authors wish to thank Antonia Ciucasel for aiding with blind behavior analysis, to Gordon Reid for providing a part of the TRPM8 clones used in the study, and to Livia Petrescu for helpful discussions.

## Author contributions

T.S. designed the study and established the methodology, performed and analyzed calcium imaging experiments, the patch clamp recordings, the behavioral experiments in rats and chickens, the spraying experiments in mice, contributed with resources, and wrote the manuscript with feedback from all authors. R.A.B. performed and analyzed calcium imaging experiments, and behavioral experiments in rats and chickens. M.I.C. performed and analyzed the C-1 experiments in mice. A.M. performed and analyzed calcium imaging experiments. V.M.C. performed and analyzed behavioral experiments in rats and chickens, and performed additional data analysis. D.E.H. performed and analyzed calcium imaging experiments, and performed additional data analysis. R.M. recorded and analyzed the s.c. temperature in sprayed mice. E.T.W. contributed with resources (chemical compounds). E.C. performed and analyzed the C-1 experiments in mice, and contributed with resources. K.Z. contributed with resources and methodology for the mouse spraying experiments. A.B. performed and analyzed calcium imaging experiments in mouse DRG neurons and contributed with resources.

## Data availability

All data generated or analyzed during this study are included in the manuscript and supporting files; source data files have been provided for Figures 1, 2, 5 and 6.

