## Supplementary Figure S1, Supplementary Figure S2, Supplementary Table S1, Supplementary movie captions for "TRPM8-dependent shaking in mammals and birds"

Supplementary figures

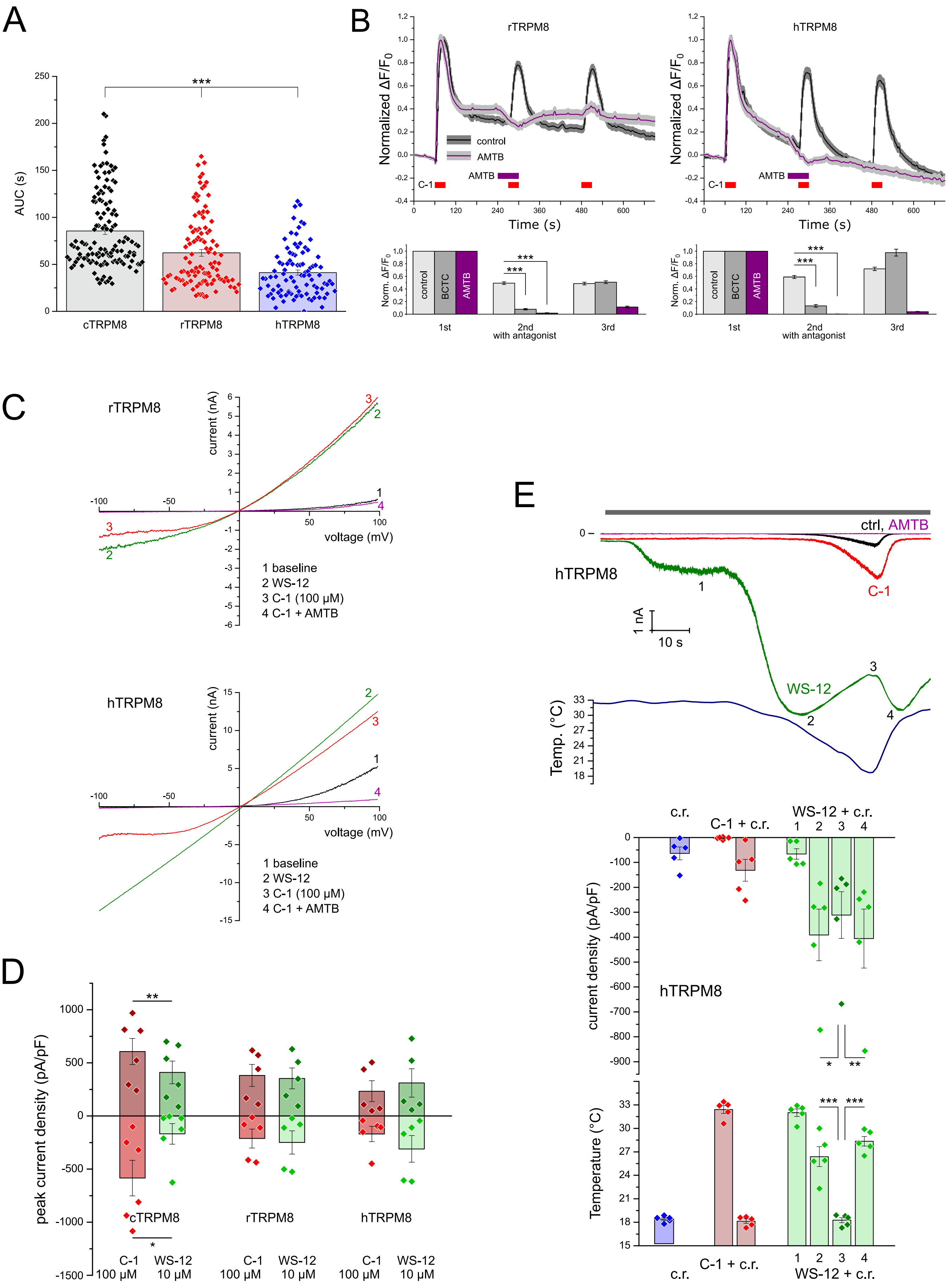

**Fig. S1. Properties of the novel agonist C-1 on recombinant TRPM8 from mammals.**
(**A**) Area under the $\Delta F/F_0$ curve of the response to 10 μM C-1 of c/r/hTRPM8 (mean ± SEM, ***, p<0.001 for each two pairs, one-way ANOVA, 129/110/93 cells for c/r/hTRPM8 respectively, same cells as in Fig. 1 B).
(**B**) The same experiment as in Fig. 1C, for r/hTRPM8. Top: from left to right: mean ± SEM traces ($\Delta F/F_0$) for control experiments with three C-1 (10 μM, 30 s each) challenges and traces from experiments where the TRPM8 antagonist AMTB (1 μM) was applied before and during the second C-1 challenge. The traces were normalized to the response at the first C-1 challenge (for hTRPM8 106/98 cells and for rTRPM8 99/94 cells from control/AMTB experiments). Bottom: from left to right, the normalized means of maximal $\Delta F/F_0$ from the above control and AMTB experiments. A similar set of experiments investigated the inhibition with BCTC (10 μM), a different TRPM8 antagonist (***, $p<0.001$, Student's *t*-test, paired; 106/99/100 cells for rTRPM8; 99/101/100 cells for hTRPM8 from control/AMTB/BCTC experiments).
(**C**) The same experiment as in Fig. 1F with representative examples of I-V relationships for the r/hTRPM8 expressed in HEK293T cells when superfused with WS-12 (10 μM), C1 (100 μM) and C1 plus AMTB (1 μM). The temperature was kept at 25 °C.
(**D**) Outward and inward peak current densities showing the species-dependent sensitivities to C-1 (100 μM) and WS-12 (10 μM), with C-1 eliciting larger currents than WS-12 in cells transfected with cTRPM8, while for hTRPM8 the reverse tendency is recorded (**, $p<0.01$; *, $p<0.05$, Student's *t*-test, paired, n=6 for cTRPM8 and n=5 for r/ hTRPM8).
(**E**) The same experiment as in Fig. 1H, performed for hTRPM8. Top: Representative inward currents recorded during cooling ramps alone (from 32-33 to 18 °C) and with agonists/antagonist exposure. All currents were measured from basal current; for C-1 and WS-12 the first column shows the current elicited by the agonist alone, before the cooling ramp was initiated and are marked with '1'. The currents recorded while the cooling ramp and a potent agonist were simultaneously applied show a characteristic dip close to minimum temperature, as seen in hTRPM8 with WS-12. The current dip is marked with '3', between the two maxima, '2' and '4'. AMTB (1 μM) abolished the currents evoked by cooling in hTRPM8. Bottom: data pooled from experiments as above, showing the average current densities and temperatures at which the currents were recorded. C-1 with cooling evoked smaller currents than WS-12 in hTRPM8 (***, $p<0.001$; **, $p<0.01$; *, $p<0.05$; one-way repeated measures ANOVA, n=5). All columns represent means ± SEM.

Rat DRG neurons

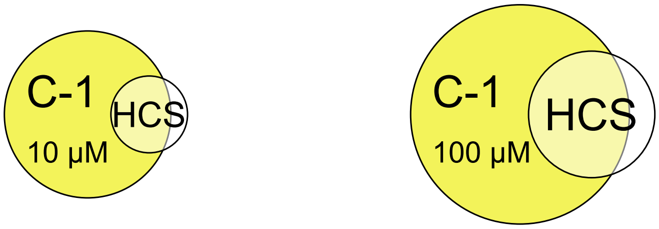


**Fig. S2. Cold-sensitive neurons activated by C-1.**
From the total 1426 neurons imaged in both experiments (with 10 and 100 µM C-1, the same shown in Fig. 2), 220 neurons (15.43%) responded to the mild cooling step (32 to 25 °C) with small amplitude calcium transients (average Δ F/F0=16.24% ± 0.02 %). These highly cold-sensitive (HCS) neurons displayed C-1 and WS-12 sensitivities: in the 10 µ M C-1 experiment, 13 of 16 HCS neurons were activated by both agonists, whereas at 100 µM C-1, 28 of 36 the HCS neurons were activated by both agonists, while one HCS neuron was only activated by C-1.

### Supplementary table

**Table S1. Summary of $EC_{50}$ and $Q_{10}$ values determined from calcium imaging and whole-cell patch clamp experiments, respectively, related to Fig. 1**

| TRPM8 ortholog | C-1, calcium imaging | | | | current increase | | current decrease | |
|---|---|---|---|---|---|---|---|---|
| | 25 °C | | 32 °C | | $1/Q_{10}$** | agonist | $Q_{10}$** | agonist |
| | $EC_{50}$ (µM)* | Hill coeff.* | $EC_{50}$ (µM)* | Hill coeff.* | | | | |
| chicken | 0.25 ± 0.01 | 1.66 ± 0.04 | 1.33 ± 0.10 | 1.95 ± 0.43 | 138.84 ± 44.35 (n=5) | C-1 (10 µM) | 1.23 ± 0.06 (n=3) | C-1 (10 µM) |
| rat | 1.62 ± 0.63 | 2.89 ± 2.34 | 2.24 ± 0.69 | 3.40 ± 1.29 | 317.90 ± 129.61 (n=5) | WS-12 (10 µM) | 1.29 ± 0.14 (n=4) | WS-12 (10 µM) |
| human | 2.43 ± 1.01 | 1.07 ± 0.34 | 4.73 ± 0.43 | 1.42 ± 0.12 | 819.56 ± 557.43 (n=5) | WS-12 (10 µM) | 1.28 ± 0.07 (n=4) | WS-12 (10 µM) |

* value ± standard error; ** average ± SEM ; the number of experiments is mentioned in the table

### Supplementary movie captions

**Movie S1. C-1 evokes robust WDS in rats.**
Excerpts from the recording of a female rat i.p. injected with C-1. Recorded and displayed at 30 fps.

**Movie S2. *Trpm8-/-* do not exhibit WDS after C-1 i.p injection, in contrast to WT.**
Top (left and right): WT C57BL/6 mice; Bottom (left and right): *Trpm8-/-* mice. Recorded and displayed at 25 fps.

**Movie S3. Mechanical stimuli evoke WDS in mice.**
Examples of WDS behavior in mice after semolina powder sprinkling. Recorded at 240 fps and displayed at variable fps; sprinkling and shaking are displayed at 30 fps.

**Movie S4. Cold water spraying evokes WDS and fast dorsal fur temperature changes in mice.**
Left: color camera Movie capture from the side; Right: FLIR camera Movie capture from above, both recorded and displayed at 30 fps.

**Movie S5. C-1 injection elicits shaking behavior in chickens.**
Examples of typical head and body shakes in C-1 s.c. injected chickens. Recorded and displayed at 30 fps.

**Movie S6. C-1 injection induces corner crowding and sleep-like behavior in chickens.**
Examples of typical corner crowding and sleep-like behavior in C-1 s.c. injected chickens. Recorded and displayed at 30 fps.

**Movie S7. Water pouring evokes vigorous shaking in farmhouse adult chickens.**
Typical body and head shakes used for water removal from feathers. Recorded at 240 fps and displayed at 30 fps.

**Movie S8. Mechanical stimuli evoke shaking in adult chickens.**
Examples of body shake behavior in adult chickens after semolina powder sprinkling. Recorded at 240 fps and displayed at 30 fps.

**Movie S9. Water spraying evokes shaking behavior in chickens.**
Examples of typical head and body shakes in water sprayed chickens recorded at 30 fps.
